# Diet-derived metabolites and mucus link the gut microbiome to fever after cytotoxic cancer treatment

**DOI:** 10.1101/2021.09.18.460647

**Authors:** Zaker Schwabkey, Diana H. Wiesnoski, Chia-Chi Chang, Wen-Bin Tsai, Dung Pham, Saira S. Ahmed, Tomo Hayase, Miriam R. Ortega Turrubiates, Rawan K. El-Himri, Christopher A. Sanchez, Eiko Hayase, Annette C. Frenk Oquendo, Takahiko Miyama, Taylor M. Halsey, Brooke E. Heckel, Alexandria N. Brown, Yimei Jin, Philip L. Lorenzi, Marc O. Warmoes, Lin Tan, Alton G. Swennes, Vanessa B. Jensen, Christine B. Peterson, Kim-Anh Do, Liangliang Zhang, Yushu Shi, Yinghong Wang, Jessica R. Galloway-Pena, Pablo C. Okhuysen, Carrie R. Daniel-MacDougall, Yusuke Shono, Marina Burgos da Silva, Jonathan U. Peled, Marcel R.M. van den Brink, Nadim Ajami, Jennifer A. Wargo, Gabriela Rondon, Samer A. Srour, Rohtesh S. Mehta, Amin M. Alousi, Elizabeth J. Shpall, Richard E. Champlin, Samuel A. Shelburne, Jeffrey J. Molldrem, Mohamed A. Jamal, Jennifer L. Karmouch, Robert R. Jenq

## Abstract

Not all cancer patients with severe neutropenia develop fever, and the fecal microbiome may play a role. In neutropenic hematopoietic cell transplant patients (n=119), 63 (53%) developed a subsequent fever and had increased fecal *Akkermansia muciniphila*, a mucus-degrading bacteria (p=0.006, corrected for multiple comparisons).

In mouse models, two therapies, irradiation and melphalan, similarly expanded *A. muciniphila*. Dietary restriction of unirradiated mice also expanded *A. muciniphila* and thinned the colonic mucus layer. Azithromycin treatment depleted *A. muciniphila* and preserved colonic mucus.

Dietary restriction raised colonic luminal pH and reduced acetate, propionate, and butyrate. Culturing *A. muciniphila* with lower pH and increased propionate prevented utilization of mucin. Treating irradiated mice with azithromycin or propionate preserved the mucus layer, lessened hypothermia, and reduced inflammatory cytokines in the colon. These results suggest that diet, metabolites and colonic mucus link the microbiome to neutropenic fever, and could guide future microbiome-based preventive strategies.

## Introduction

One of the most common and potentially serious complications of cancer therapy is neutropenia and subsequent infectious complications, with an estimated mortality of nearly 10% (*1*) as well as 100,000 hospitalizations and over $2.7 billion in hospitalization costs annually in the United States (*2*). At particularly high risk are patients undergoing chemotherapy for hematological malignancies including acute leukemias and high-grade lymphomas or receiving hematopoietic cell transplantation (HCT) after myeloablative conditioning (*3*).

The degree and duration of neutropenia has long been identified as a critical clinical parameter predicting infection (*4*). More recently, the role of the intestinal microbiome in neutropenia-related infections has been increasingly appreciated with the majority of documented bacterial infections arising from the gastrointestinal tract (*5, 6*). Most patients, however, will not have an infectious etiology identified, and it is not well-understood why only some 50% of patients with profound neutropenia become febrile. Whether other intestinal commensal bacteria, primarily nonpathogenic bacteria, could also play a role in infection risk, has not been extensively studied. Herein we sought to gain insight into the pathophysiology of neutropenic fever using a combination of human samples and experimental mouse models.

## Results

We began with an examination for a potential relationship between the composition of intestinal bacteria and fever, in patients who developed neutropenia in the setting of HCT. We examined a cohort of 119 patients at our center who all developed neutropenia following HCT conditioning. Of these, 56 (47%) remained afebrile over the next 4 days, while 63 (53%) developed a fever, of which 7 were found to have a bloodstream infection, including 5 with Enterobacteriaceae and 2 with oral streptococci. In the microbiome analyses described below, patients with bloodstream infections were included with those who developed fever, though we also analyzed excluding these patients and found very similar results. Prior to stool collection, patients were treated with prophylactic levofloxacin given daily per our institutional standard. Additional patient characteristics are provided in Table 1. Receiving an autologous HCT, or an HCT following a preparative regimen based on both busulfan and melphalan, were associated with increased fever, likely reflecting increased conditioning intensity.

**Table 1.**
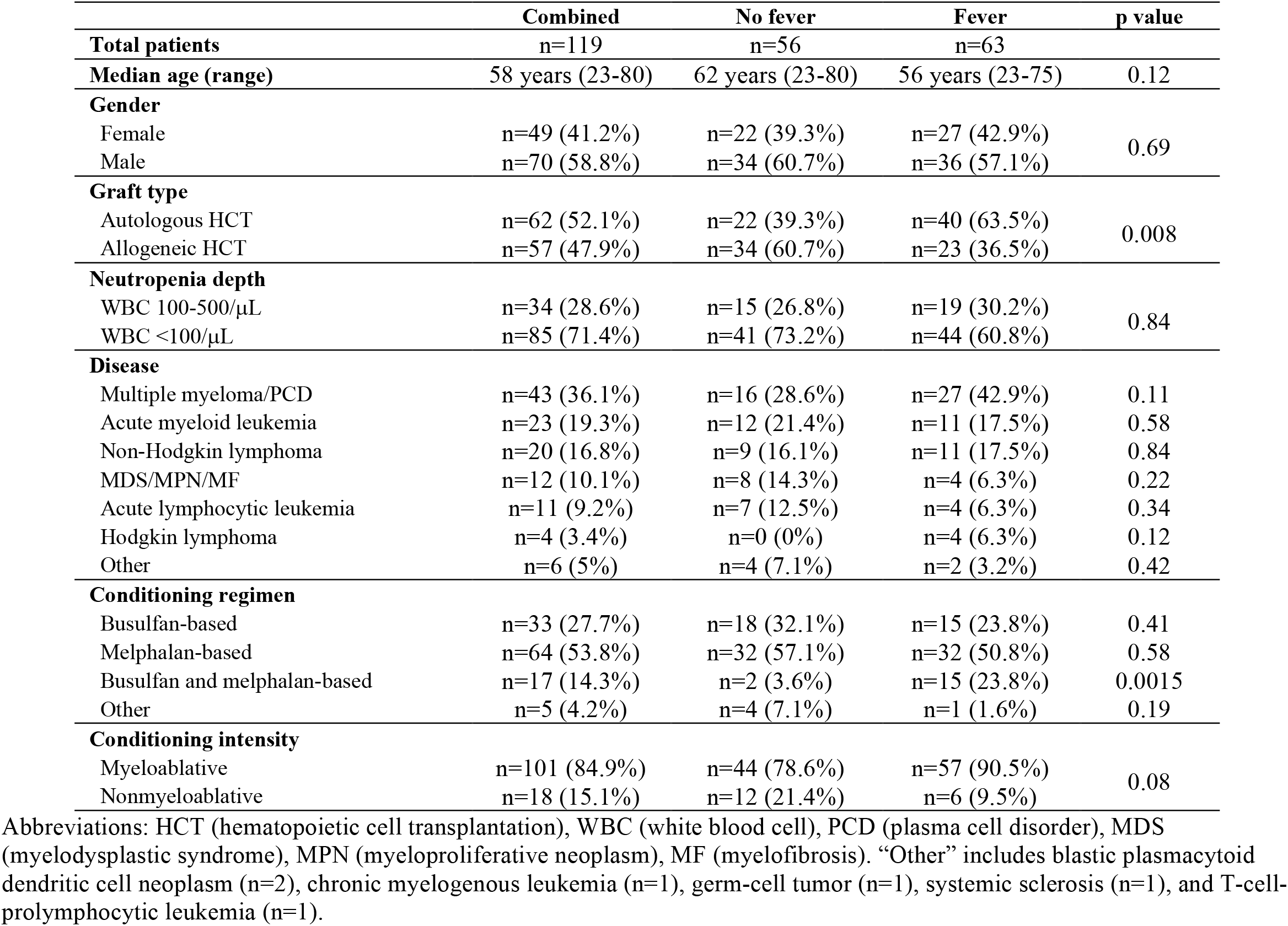
Patient characteristics

When analyzing stool samples collected at the onset of neutropenia, we found evidence for microbiome differences between those who did or did not develop fever over the next 4 days. There was a significant difference in beta-diversity (p=0.02, permutational MANOVA, Figure 1A). Patients who later developed fever had increased relative abundance of bacteria from the genus *Akkermansia* (p=0.006, adjusted for multiple comparisons), as well as bacteria from the genus *Bacteroides* (p=0.01), while bacteria from the class Bacilli and from the order Erysipelotrichales were increased in patients that were afebrile (Figures 1B, C and D). Notably, bacterial taxa associated with bloodstream infections, included Enterobacteriales, *Streptococcus* and *Enterococcus (7)* were not associated with fever.

**Figure 1.**
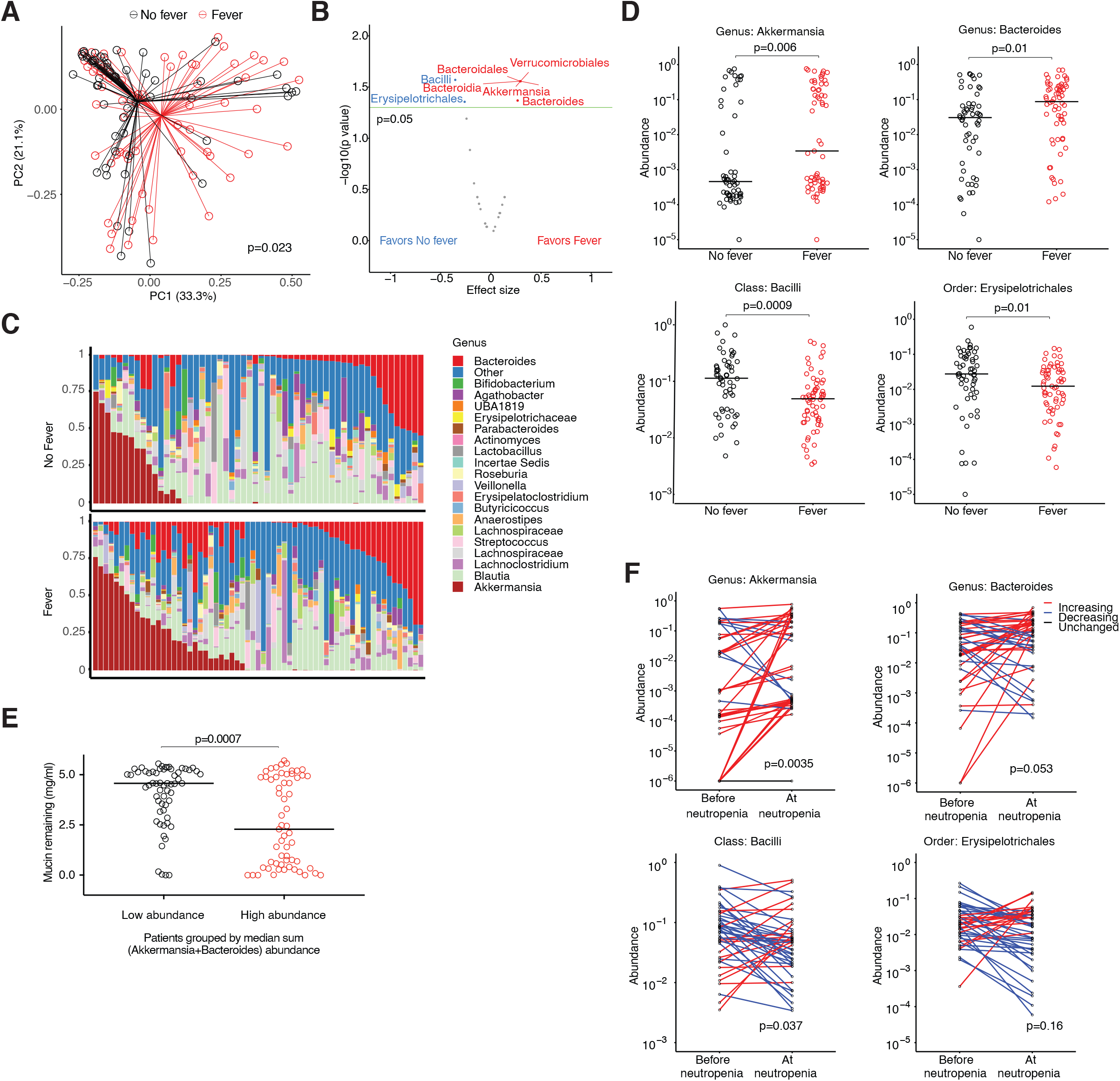
Mucus-degrading intestinal bacteria are associated with development of fever following onset of neutropenia in HCT patients. Intestinal microbiome parameters at neutropenia onset and subsequent fever were evaluated in a cohort of patients undergoing HCT. Stool samples were collected at onset of neutropenia (+/- 2 days), and fever outcome was determined by inpatient monitoring every 4 hours in the subsequent 4 days after collection. A) Following 16S rRNA gene sequencing, Principal Coordinates Analysis (PCoA) was performed on weighted UniFrac distances. Statistical significance was determined by Permutational Multivariate Analysis Of Variance (PERMANOVA) testing. B) Volcano plot of bacterial taxa that were differentially abundant in A). Taxa above the green line have a p value less than 0.05; p values were adjusted for multiple comparisons. C) Relative abundances of bacteria at the genus level in samples from A) are indicated in stacked bar graphs. D) Relative abundances of bacteria of the indicated taxa are depicted for samples from A); p values were adjusted for multiple comparisons. E) Mucin glycan consumption by frozen aliquots of stool samples in A) was assayed. Fecal bacteria were cultivated in liquid media supplemented with porcine gastric mucin as the predominant source of carbon, followed by quantification of remaining mucin glycans after 48 hours. Samples were stratified by median sum abundance of *Akkermansia* and *Bacteroides*. F) In the subset of patients who later developed neutropenic fever, relative abundances of bacteria from the indicated taxa in stool samples collected at onset of neutropenia were compared to results of a baseline stool sample collected earlier in the hospitalization, using the Wilcoxon signed-rank test.

The genus *Akkermansia* currently includes only one species, *Akkermansia muciniphila (A. muciniphila)*, while *Bacteroides* is quite diverse. Interestingly, *A. muciniphila* and several members of *Bacteroides* are known to have mucus-degrading capabilities (*8*), and so we asked if intestinal bacteria from patients with febrile neutropenia had an increased ability to degrade mucin glycans. To evaluate this we adopted an approach where certain carbohydrates including mucin glycans can be quantified from liquid samples using periodic acid-Schiff’s reagent (Supplementary Figure 1A) (*9*). We found that such a method could quantify the concentration of glycans derived from commercially available porcine gastric mucin (PGM) in media. Following a 48-hour culture we could quantify a reduction in glycan concentration in media inoculated with isolates of *A. muciniphila* (ATCC BAA-835) and *Bacteroides thetaiotaomicron* (*B. thetaiotaomicron*, ATCC 29148), but glycan levels were not altered by a non-mucolytic isolate of *E. coli* (ATCC 700926, Supplementary Figure 1B). We applied this assay to aliquots of patient samples from Figure 1, and found that samples from patients with higher combined abundances of *Akkermansia* and *Bacteroides* were more effective at consuming mucin glycans (Figure 1E).

We then asked if abundances of bacteria at onset of neutropenia were changed from baseline in HCT patients. In a majority of patients from our cohort (32 of 56 without fever and 44 of 63 with fever), baseline stool samples had been collected at least 4 days prior to onset of neutropenia and were available for comparison. Interestingly, we found that patients who later developed fever had significantly increased *Akkermansia* and reduced Bacilli at onset of neutropenia compared to baseline, while afebrile patients had no significant changes over time in any of the bacteria associated with fever (Figure 1F, Supplementary Figure 1C). In summary, our evaluation of the microbiome in the setting of clinical neutropenia onset identified an increase in the abundance of bacteria with mucolytic properties in patients who later developed fever, and *Akkermansia* in particular was significantly increased from baseline in patients who later developed fever.

The fact that our previous analyses had been performed only in patients not receiving broad-spectrum antibiotics suggested a non-antimicrobial mechanism mediating the dysbiotic expansion of mucolytic bacteria. Thus, we next sought to test the hypothesis that transplant conditioning could change the composition of the intestinal microbiome using a murine model which permitted experimental evaluation of the impact of cytotoxic therapy. We began with total body radiotherapy. C57BL/6 mice were exposed to a single myeloablative dose of total body radiotherapy (9 Gy RT), and their stool samples were evaluated 6 days later by 16S rRNA gene sequencing. This time point was chosen because some mice would often become moribund by day 7. Strikingly, the microbiome of mice on day 6 was markedly changed compared to unarriadated mice (Figure 2A). Moreover, the profile was highly reminiscent of that seen in HCT patients with febrile neutropenia, showing increases in the abundance of *Akkermansia*, and to a lesser degree, *Bacteroides* (Figure 2B-D). Notably, we did not observe compensatory reductions in Bacilli or Erysipelotrichales, which had been seen in patients with febrile neutropenia. Rather, we found reductions in bacteria from the family Muribaculaceae, a recently named group of bacteria which is commonly found in high abundance in mice but is usually a minor contributor in the intestinal tract of humans (*10*). We asked if this change in bacterial composition was accompanied by functional changes, and found that bacteria derived from stool samples of irradiated mice more efficiently degraded mucin glycans than bacteria from unirradiated mice in vitro (Figure 2E). We then asked if the dense colonic mucus layer, which normally separates bacteria-rich luminal contents from the colonic epithelium, was affected by irradiation in mice. To evaluate this, colonic tissue samples were cut cross-sectionally and mucus layer thickness was averaged across 8 equally-spaced circumferential sites (Supplementary Figure 2). Using this systematic approach, we found that the mucus layer was significantly thinner in irradiated mice compared to normal mice (Figure 2F).

**Figure 2.**
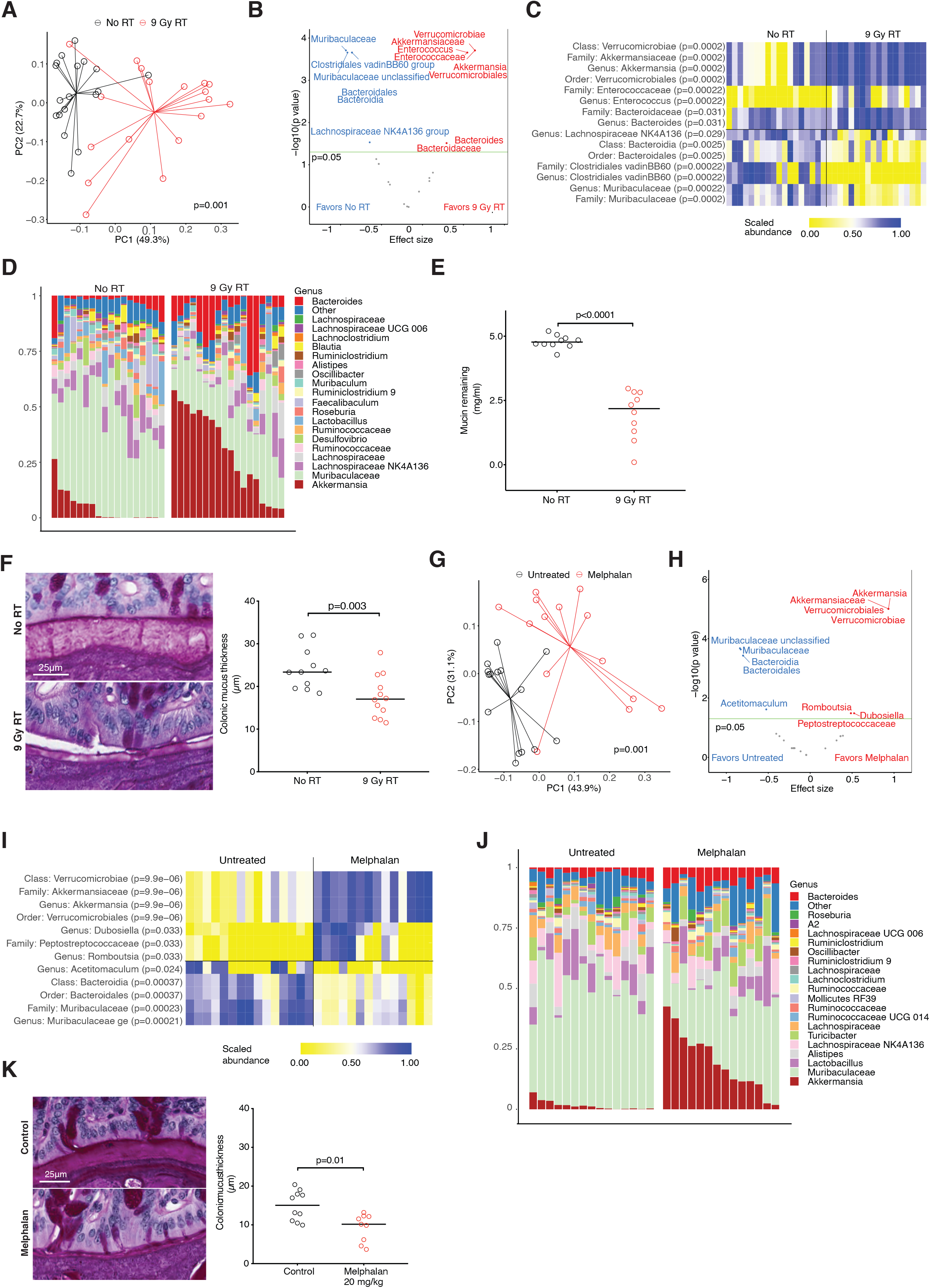
Systemic cytotoxic therapy increases the relative abundance of mucus-degrading intestinal bacteria in mice. Evaluation of intestinal microbiome parameters was performed in adult C57BL/6 female mice 6 days following total body radiotherapy (9 Gy RT, panels A-E) or 6 days following melphalan therapy (20 mg/kg, panels G-K). A) After 9 Gy RT, PCoA was performed on weighted UniFrac distances; combined results of 3 experiments. B) Volcano plot of bacterial taxa that were differentially abundant in A); p values were adjusted for multiple comparisons. C) Heat map of scaled relative bacterial abundances of the indicated taxa are depicted for samples from A). D) Relative abundances of bacteria at the genus level in samples from A) are indicated in stacked bar graphs. E) Bacteria from frozen stool samples collected from mice in A) were evaluated for mucin glycan consumption; combined results of 2 experiments. F) Thickness of the dense inner colonic mucus layer was evaluated histologically in mice in A). Representative images are provided with combined results of 3 experiments. G) After melphalan therapy, PCoA was performed on weighted UniFrac distances; combined results of 3 experiments. H) Volcano plot of bacterial taxa that were differentially abundant in G); p values were adjusted for multiple comparisons. I) Heat map of scaled relative bacterial abundances of the indicated taxa are depicted for samples from G). J) Relative abundances of bacteria at the genus level in samples from G) are indicated in stacked bar graphs. K) Thickness of the dense inner colonic mucus layer was evaluated histologically in mice in G). Representative images are provided with combined results of 2 experiments.

Myeloablative RT, previously a foundational pillar which made HCT possible, has been progressively replaced by chemotherapy, particularly alkylating agents (*11*). We thus treated mice with the alkylating agent melphalan, and found that treatment led to significant changes in the microbiome, marked particularly by an increase in the abundance of *Akkermansia*, similar to that seen after RT (Figures 2G-J). An expansion of *Bacteroides* was seen, though this was not statistically significant after correction for multiple comparisons, while a loss of Muribaculaceae, similar to that following RT, was also seen. Histological analysis demonstrated that the mucus layer was significantly thinner in melphalan-treated mice, similar to RT (Figure 2K).

We asked why the intestinal microbiome appeared to be impacted similarly in response to different cytotoxic therapies. We first evaluated for direct effects of RT on intestinal bacterial composition by irradiating mouse fecal pellets and cultivating bacteria on agar plates. We then swabbed all bacterial colonies that grew and evaluated their taxonomical composition by 16S rRNA gene sequencing. We found that exposure to irradiation resulted in no enrichment for *Akkermansia* or *Bacteroides*, though *Akkermansia* abundance was low due to its relatively slow growth rate in vitro (Supplementary Figure 3A). We also introduced bacteria from irradiated fecal pellets orally to mice previously treated with an oral decontaminating antibiotic cocktail. We again found that exposure to irradiation had no discernible effect on the composition of bacterial populations, including no enrichment for *Akkermansia* or *Bacteroides* (Supplementary Figure 3B).

Diet is known to be a major determinant of intestinal microbiome composition. We asked if RT could be impacting the microbiome composition indirectly, by causing a reduction in intake of food in mice. We individually housed mice in metabolic cages following RT to quantify effects on food and water intake. We found that within 2 days following RT, mice had reduced their oral intake to approximately 2 grams a day, or a 50% reduction (Figure 3A). To evaluate whether dietary restriction (DR) could impact intestinal microbiome composition and the colonic mucus layer, we took normal, unirradiated mice, and limited their access of chow to 2 grams per mouse per day for 7 days. We found that dietary restriction resulted in marked changes in the microbiome characterized primarily by expansion of *Akkermansia* and, to a lesser extent, an expansion of *Bacteroides* and loss of Bacilli (Figures 3B-E). Mucin glycan degradation was more robust after DR (Figure 3F), and the colonic mucus layer was also significantly thinner (Figure 3G). To evaluate if DR was impacting mucin production, we evaluated goblet cells, specialized epithelial cells that are the primary producers of mucin in the colon. Neither the numbers of goblet cells per crypt, nor the combined surface area of goblet cells in a cross section of colonic tissue, were impacted by DR (Supplementary Figure 4A). RNA expression of the gene encoding the predominant mucin in the small and large intestine, Muc2, was not significantly changed in colonic tissue homogenates from mice following DR (Supplementary Figure 4B). Altogether, these results indicated that a reduction in oral nutrition was sufficient to produce a thinner colonic mucus layer, possibly through an increase in mucin-degradation leading to increased consumption of mucin, while the production of mucin appeared to be intact.

**Figure 3.**
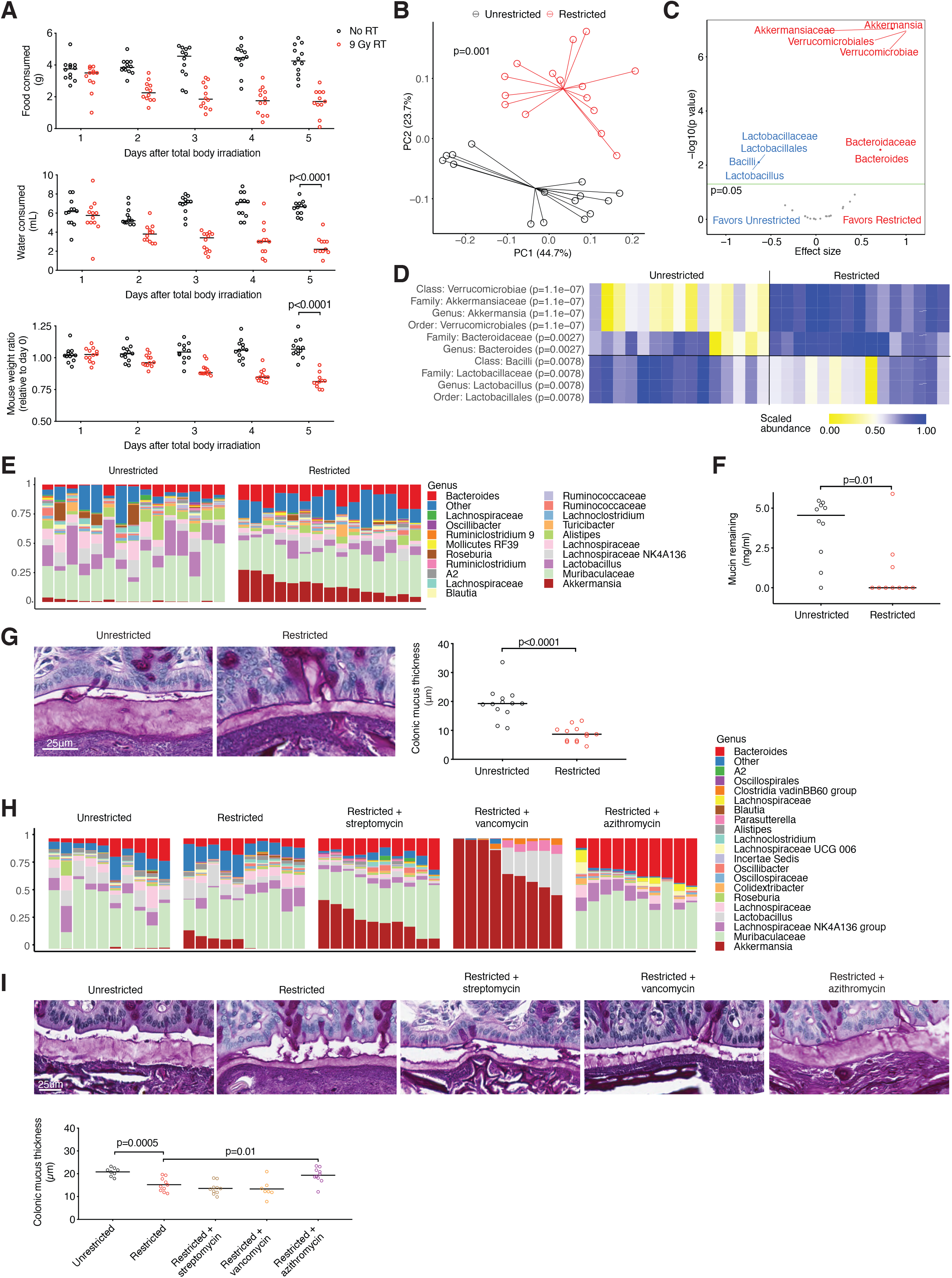
Dietary restriction increases the relative abundance of mucus-degrading intestinal bacteria in mice. A) After 9 Gy RT, mice were individually housed in metabolic cages and monitored daily for food consumption, water consumption, and weight. B) Intestinal microbiome parameters were evaluated in normal mice after undergoing dietary restriction (2 g/mouse/day) for one week. PCoA was performed on weighted UniFrac distances; combined results of 3 experiments. C) Volcano plot of bacterial taxa that were differentially abundant in B); p values were adjusted for multiple comparisons. D) Heat map of scaled relative bacterial abundances of the indicated taxa are depicted for samples from B). E) Relative abundances of bacteria at the genus level in samples from A) are indicated in stacked bar graphs. F) Bacteria from frozen stool samples collected from mice in B) were evaluated for mucin glycan consumption; combined results of 2 experiments. G) Thickness of the dense inner colonic mucus layer was evaluated histologically in mice in B). Representative images are provided with combined results of 3 experiments. H) Mice underwent dietary restriction as in B), with the addition of narrow-spectrum antibiotics administered in the drinking water starting 5 days prior to onset of restriction. Relative abundances of bacteria at the genus level in samples are indicated in stacked bar graphs; combined results of 2 experiments. I) Thickness of the dense inner colonic mucus layer was evaluated histologically in mice in H). Representative images are provided with combined results of 2 experiments.

Commercially-available mice lacking *Akkermansia* are not easy to find (*12*). To help evaluate the specific contribution of *Akkermansia* to mucus thinning during DR, we turned to narrow-spectrum antibiotics. Specifically, we evaluated streptomycin which we found depleted certain gram-positive bacteria, vancomycin which depleted both gram-positive bacteria and some Bacteroidetes, and azithromycin, which we found depleted *Akkermansia* as well as some gram-positive populations (Figure 3H). We found that neither streptomycin nor vancomycin had a significant effect on mucus barrier loss, while azithromycin treatment was able to prevent thinning of the colonic mucus layer (Figure 3I), indicating that *Akkermansia* could be required for increased mucolysis during dietary restriction. To identify mechanistic links between diet and microbiome composition, we hypothesized that reduced oral dietary intake was perturbing normal commensal bacterial metabolism in the intestinal lumen, and began by asking if metabolic substrates entering the colon were impacted by dietary restriction. To evalute this we performed bomb calorimetry on cecal contents of mice and found that restricted mice had less calories entering the colon (Figure 4A). Among the most abundant products of intestinal bacterial metabolism are organic acids, thus we quanitified pH in the colonic lumen, and found that dietary restriction resulted in a raised pH, indicating overall reduced metabolism (Figure 4B). To better characterize this rise in pH, we quantified specific bacterial metabolites using ion-chromatography mass spectometry, and found that dietary restriction led to reduced concentrations of acetate, propionate and butyrate, and, interestingly, increased succinate (normalized results in Figure 4C, raw results in Supplementary Figure 5A). Succinate is a metabolic precursor of propionate (*13*), suggesting that dietary restriction could be inhibiting enzymatic conversion of succinate to propionate.

**Figure 4.**
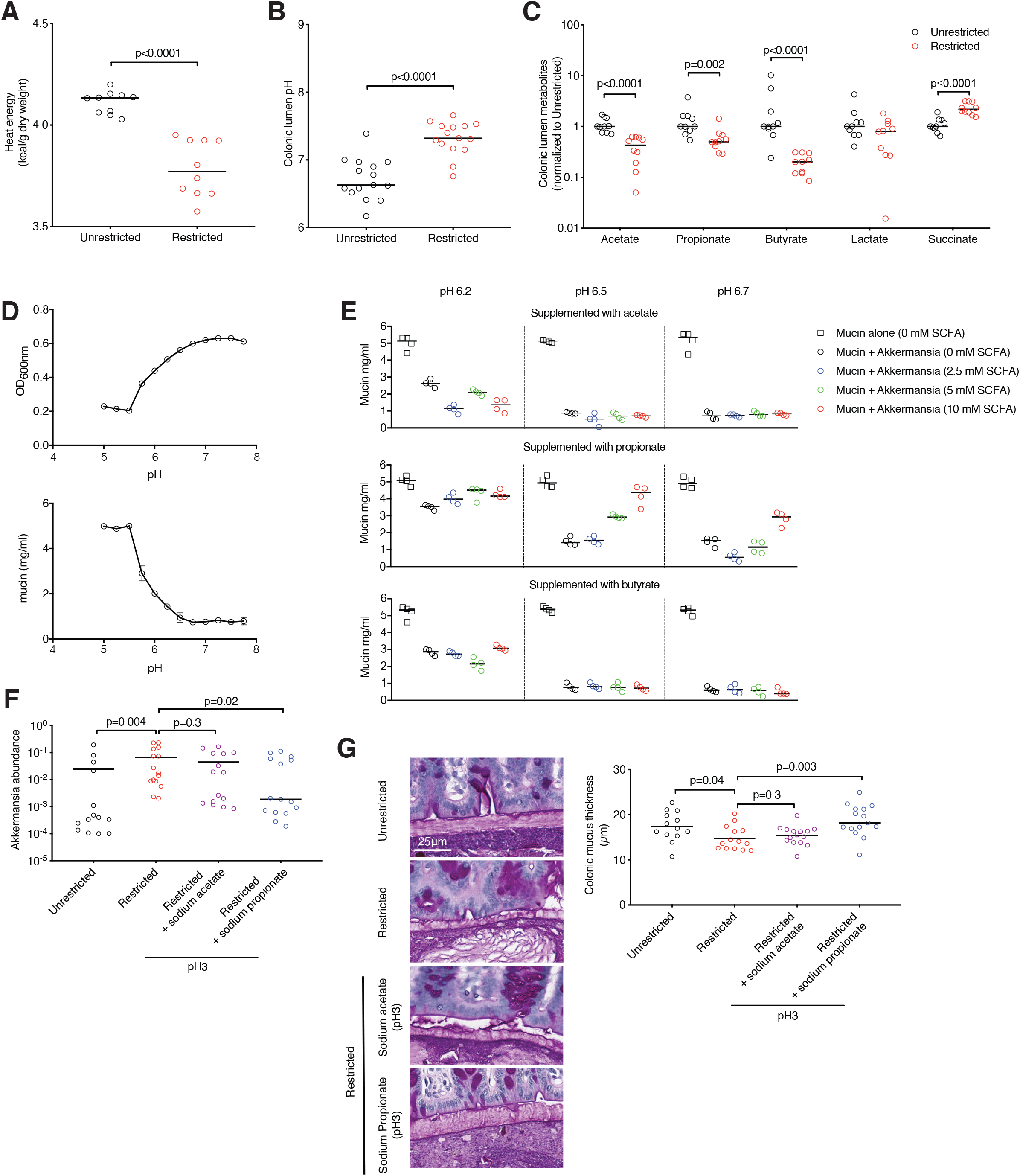
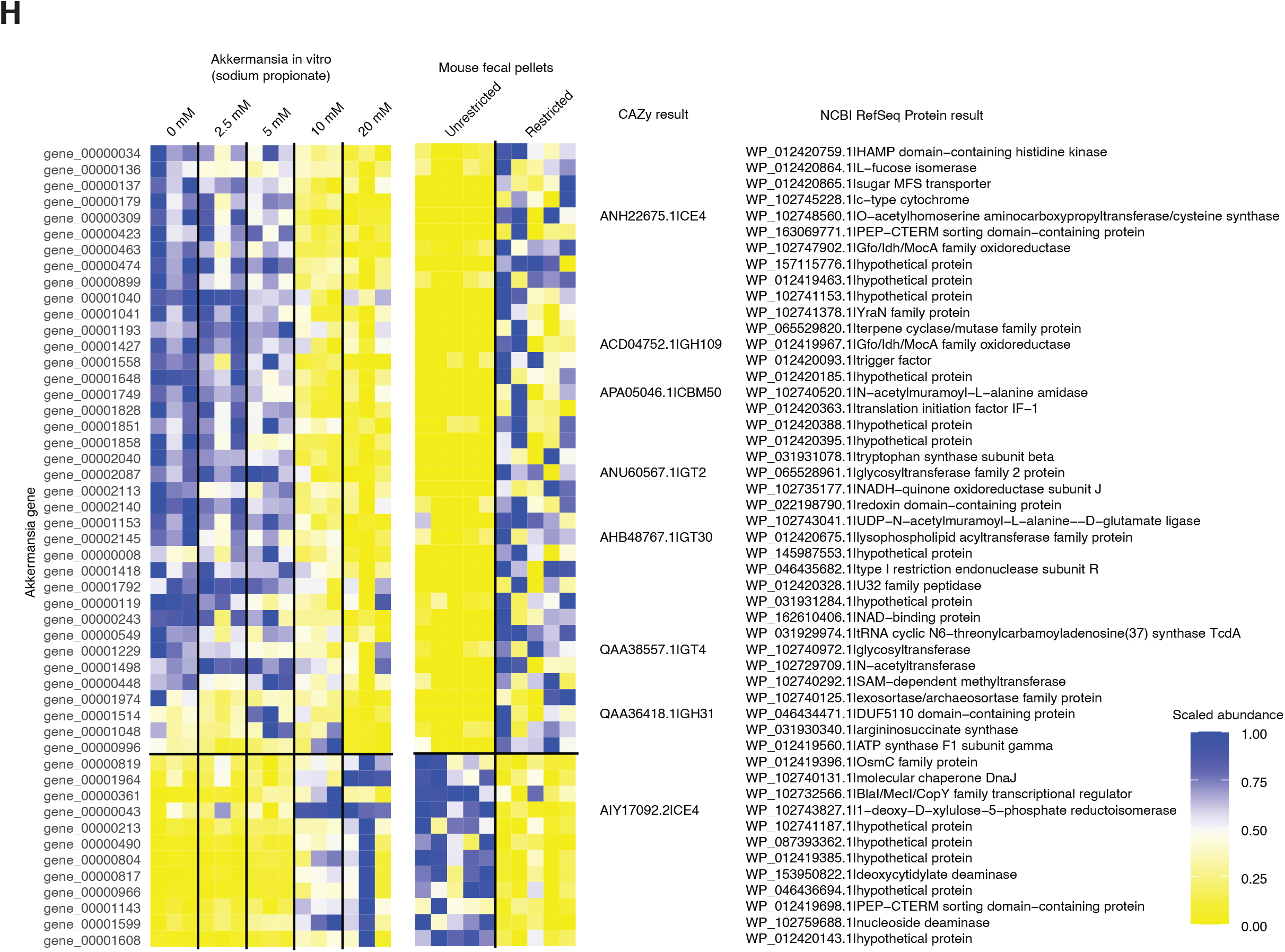
Bacterial metabolites link dietary restriction to mucolytic bacteria. A) In mice that underwent one week of dietary restriction, cecal luminal contents were assessed for caloric content by bomb calorimetry; combined results of 2 experiments. B) Colonic luminal contents were assessed for pH in mice following one week of dietary restriction; combined results of 3 experiments. C) Metabolites from samples in B) were quantified using ion chromatography-mass spectrometry (IC-MS); combined results of 2 experiments. D) A murine isolate of *A. muciniphila* (MDA-JAX001) was cultivated under anaerobic conditions of varying pH in 4 replicates, and growth and mucin glycan consumption were quantified after 48 hours of culture; results of one of two experiments with similar results. E) *A. muciniphila* (MDA-JAX001) was cultivated under varying pH and varying concentrations of sodium acetate, sodium propionate, and sodium butyrate in 4 replicates, and mucin glycan consumption was quantified after 48 hours of culture; results of one of two experiments with similar results. F) Normal mice received one week of dietary restriction, as well as supplementation with sodium acetate or sodium propionate in the drinking water, acidified to pH3. Relative abundances of *Akkermansia* was quantified by 16S rRNA gene sequencing; combined results of 3 experiments. G) Thickness of the dense inner colonic mucus layer was evaluated histologically in mice in F). Representative images are provided with combined results of 3 experiments. H) Transcriptomic profiling identifies *A. muciniphila* genes similarly regulated by diet in vivo and propionate in vitro. RNA sequencing was performed on *A. muciniphila* (MDA-JAX001) cultivated at pH 6.5 with varying concentrations of sodium propionate as in E) in 3 replicates, and on fecal pellets from mice following one week of dietary restriction (n=5). Sequences aligning with the genome of *A. muciniphila* (MDA-JAX001) were quantified, and the scaled abundances of the subset of genes similarly regulated by diet and propionate are depicted in the heat map, along with annotations obtained using both the CAZy and NCBI RefSeq Protein databases.

We asked if DR-induced metabolic changes in the colonic lumen could be impacting on *A. muciniphila* behavior. To study this, we turned to our in vitro mucin glycan consumption assay. We isolated a novel *A. muciniphila* (MDA-JAX001) strain from the feces of C57BL6 mice, introduced it to liquid media supplemented with PGM, and evaluated the effects of varying pH either alone or combined with the presence of physiological concentrations of acetate, propionate, and butyrate. Interestingly, we found that progressively lowering the pH conditions of bacterial media below 7 led to increased inhibition of *A. muciniphila* in terms of both growth and mucin glycan degradation (Figure 4D). We also found that higher levels of propionate had inhibitory effects on mucin glycan utilization by *A. muciniphila* (Figure 4E) and also led to delays in growth (Supplementary Figure 5B) while acetate and butyrate each had negligible effects on *A. muciniphila* behavior. To see if a combination of acidity and propionate could also suppress *A. muciniphila* in vivo, we evaluated supplementing mice during DR with acidified sodium propionate in the drinking water, and found that this reduced fecal pH (Supplementary Figure 5C), mitigated expansion of *Akkermansia* (Figure 4F) and prevented thinning of the mucus layer (Figure 4G). Interestingly, similar treatment with acidified sodium acetate, despite lowering the pH in the colonic lumen, had no such preventative effect (Figures 4F-G), while drinking water with sodium propionate at neutral pH was sufficient to prevent mucus thinning and trended towards preventing *Akkermansia* expansion (Supplementary Figure 5D-E). Altogether, these results indicate that a reduced level of propionate following DR and a higher pH together support increased mucolytic activity.

To explore how DR and propionate can modulate growth and mucin utilization by *A. muciniphila*, we profiled the *A. muciniphila* transcriptome under various conditions in vivo and in vitro. We began with determining the circularized genomic sequence of our murine *A. muciniphila* isolate (MDA-JAX001), and identified 1935 putative proteins (Supplementary Figure 6A). We then performed RNA sequencing on stool samples from mice, as well as *A. muciniphila* in vitro. We identified 186 genes modulated by DR in vivo (Mann-Whitney U test unadjusted < 0.05), and 392 genes modulated by propionate in vitro (Kruskal-Wallis unadjusted < 0.05). Evaluating the correlation of effect sizes in these two settings, we found that propionate-related effects only explained changes seen in DR at a proportion of 0.05, though the slope was significantly non-zero (p<0.0001, Supplementary Figure 6B). We identified 50 genes that concordantly changed in vivo during DR and in vitro in low concentrations of propionate (Figure 4H), and representations of gene expression with respect to the genome are depicted in Supplementary Figure 6C). Mucins are glycoproteins predominately capped by fucose and sialic acid residues at their branching terminals. Notably, upregulated genes in the settings of DR and low propionate included L-fucose isomerase which interconverts fucose and fucolose, as well as a member of the glycosyl hydrolase enzyme family 109, which have been shown to cleave oligosaccharide chains on glycoproteins and glycolipids found on the surface of erythrocytes that determine ABO blood types (*14*). Also upregulated is a member of the Idh/MocA family of oxidoreductases, which can play a part in sialic acid utilization (*15*). In contrast, some of the genes upregulated in the settings of an unrestricted diet or high propionate include enzymes that are critical in producing nucleotides, such as deoxycytidylate deaminase and nucleoside deaminase, indicating a relative downregulation of carbohydrate utilization genes relative to housekeeping functions such as synthesizing DNA and RNA components.

Finally, we asked if either of the two strategies that we had identified as effective in preventing loss of colonic mucin during dietary restriction, azithromycin or propionate, could also reduce systemic inflammation following cytotoxic therapy. To evaluate this, we returned to our total body RT model and tested the addition of azithromycin or propionate. We found that each strategy was effective in preserving colonic mucus layer thickness (Figure 5A). Additionally, we non-invasively measured ocular surface temperatures in mice adopting a published method (*16*). In contrast to humans, mice are known to develop hypothermia in response to exposure to inflammatory ligands such as LPS (*17*) and in models of sepsis (*18*), an observation attributed possibly to differences in body size. We found that total body radiotherapy indeed produced significant hypothermia detectable as early as 1 day following RT, with temperatures of irradiated mice further decreasing over the next several days, while the temperatures of irradiated mice supplemented with azithromycin or propionate were less depressed at day 6 (Figure 5B). These results suggested that strategies to inhibit mucolytic activity of *A. muciniphila* were effective at reducing inflammation in irradiated mice. To explore if this was indeed the case, we directly characterized the degree of inflammation in colonic tissue by quantifying levels of a panel of cytokines (Figure 5C). We found that IL-1β, CCL2, CCL7, IL-22, CXCL1, and CXCL10 were all elevated in colonic tissues of mice following RT, but were reduced with the addition of azithromycin treatment. Propionate treatment also prevented elevation of all of these cytokines, with the exception of CXCL1 which remained elevated. We did not observe elevations of TNF following RT, nor effects of azithromycin or propionate on TNF levels. Corroborating a less complete suppression of colonic inflammation by propionate, we found that azithromycin was very effective at preventing outgrowth of *Akkermansia* in mice following RT, while propionate was not effective (Figure 5D). Altogether, results from interventional experiments in mice following RT indicated that azithromycin therapy was highly effective at eliminating intestinal *Akkermansia*, preserving colonic mucus, and preventing colonic inflammation and hypothermia, while propionate therapy was less effective but nevertheless significantly prevented colonic mucus thinning and hypothermia, and largely abrogated much of the colonic inflammation that occurred after RT.

**Figure 5.**
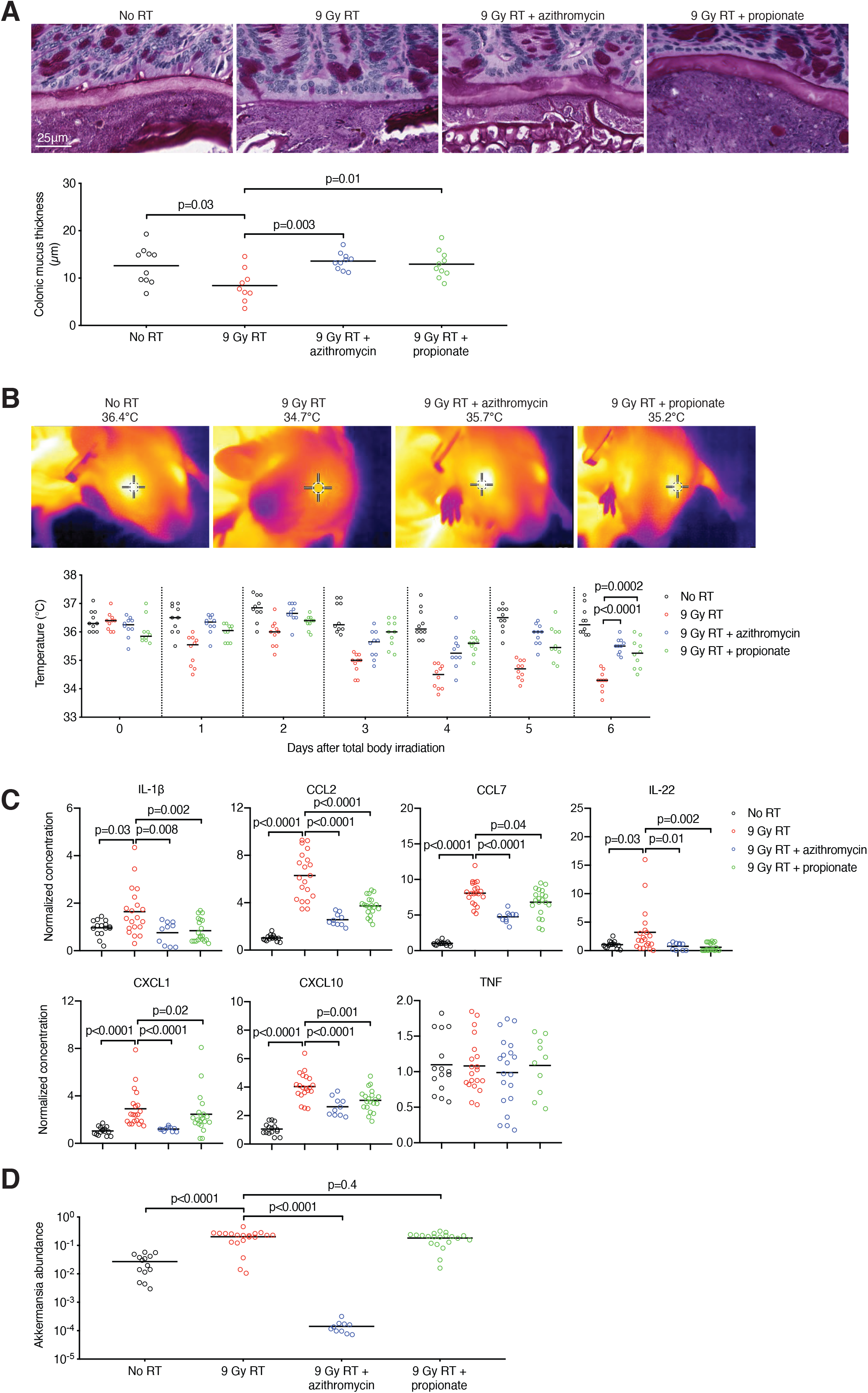
Strategies targeting mucolytic bacteria in mice receiving RT preserve colonic mucus, reduce hypothermia, and reduce colonic inflammation. In the setting of 9Gy RT, mice were treated with azithromycin or sodium propionate. A) Thickness of the dense inner colonic mucus layer was evaluated histologically. Representative images are provided with combined results of 2 experiments. B) Ocular temperatures were monitored daily. Representative images 6 days after RT are provided with combined results of 2 experiments. C) On day 6 after RT, mice were harvested and colonic tissues was processed to quantify levels of cytokines. Combined results of 3 experiments. D) Relative abundances of *Akkermansia* on day 6 after RT was quantified by 16S rRNA gene sequencing. Combined results of 3 experiments.

## Discussion

In our study, we found that bacteria with mucus-degrading capabilities, especially *Akkermansia*, were enriched in neutropenic patients who later developed fever. This is, to our knowledge, the first report of mucus-degrading intestinal bacteria being associated with this common complication of cancer therapy. Mucus degraders, however, have been previously implicated in other disease settings, including inflammatory bowel disease (*19, 20*), graft-versus-host disease (*21*) and colonic epithelial carcinogenesis (*22*). *A. muciniphila*, identified in 2004 as a specialized intestinal mucin-degrading commensal (*23*), has in particular been associated with increased colitis (*24*) and colonic graft-versus-host disease (*21*) in mouse models. Other mucin degraders include members of the genus *Bacteroides*, which have been associated with murine colitis (*25*) and are particularly well-studied due to the availability of methods to manipulate genetically tractable *Bacteroides* isolates (*26*). *Ruminococcus gnavus* is another mucin-degrading species that has been well-studied (*27*) and has been clinically associated with inflammatory bowel disease (*28*).

Interestingly, in other clinical settings, *A. muciniphila* is associated with potentially beneficial health effects. Loss of *A. muciniphila* has been observed in individuals with metabolic conditions, including obesity and Type 2 diabetes, and supplementation with *A. muciniphila* may mediate a clinical improvement (*29*). Increased *A. muciniphila* has also been associated with enhanced responses to PD1-blockade cancer immunotherapy in patients with lung and urothelial cancers, and superior tumor responses in mice (*30*).

We observed that *A. muciniphila* can increase following cytotoxic chemotherapy in some patients undergoing HCT. Radiotherapy or melphalan therapy also produced these changes in mice, and we found that this is likely driven by reductions in oral dietary intake. A link between restrictions in diet and *A. muciniphila* has been observed before, including in subjects after Islamic fasting (*31*) or following as little as three days of deliberate underfeeding in the context of a clinical trial (*32*). In mice, intermittent 16 hour fasting for one month resulted in increased *A. muciniphila* levels (*33*), as did 4 days after switching from oral to parenteral nutrition delivered following internal jugular vein catheterization (*34*), and consuming a fiber-depleted diet (*35*).

Why diet and *A. muciniphila* are closely linked is not as well-understood. We found that propionate levels in the intestinal lumen are reduced with dietary restriction, and that propionate can mediate suppressive effects on utilization of mucin glycans by *A. muciniphila*. In addition to *A. muciniphila, Staphylcoccus aureus* has also been shown to be specifically inhibited by propionate, while acetate and butyrate were not effective (*36*). Interestingly, *A. muciniphila* has been reported to produce propionate following metabolism of mucin-derived carbohydrates (*37*), and thus the presence of propionate, as a metabolic end product, may serve as a feedback mechanism to suppress excessive utilization of mucin glycans. Inhibition of *A. muciniphila* by propionate was more pronounced at lower pH settings, which could be due to better penetration of propionate through bacterial cell membranes in its protonated state, as has been observed before with acetate (*38*).

In summary, we have found that the intestinal microbiome of patients who developed fever in the setting of neutropenia was enriched in mucus-degrading bacteria. Further experimentation in mice identified interrelated aspects of diet, metabolites, and intestinal mucus. These results suggest that development of novel approaches, including dietary, metabolite, pH and antibacterial strategies, can potentially better prevent fevers in the setting of neutropenia following cancer therapy.

## Figure legends

**Figure S1.**
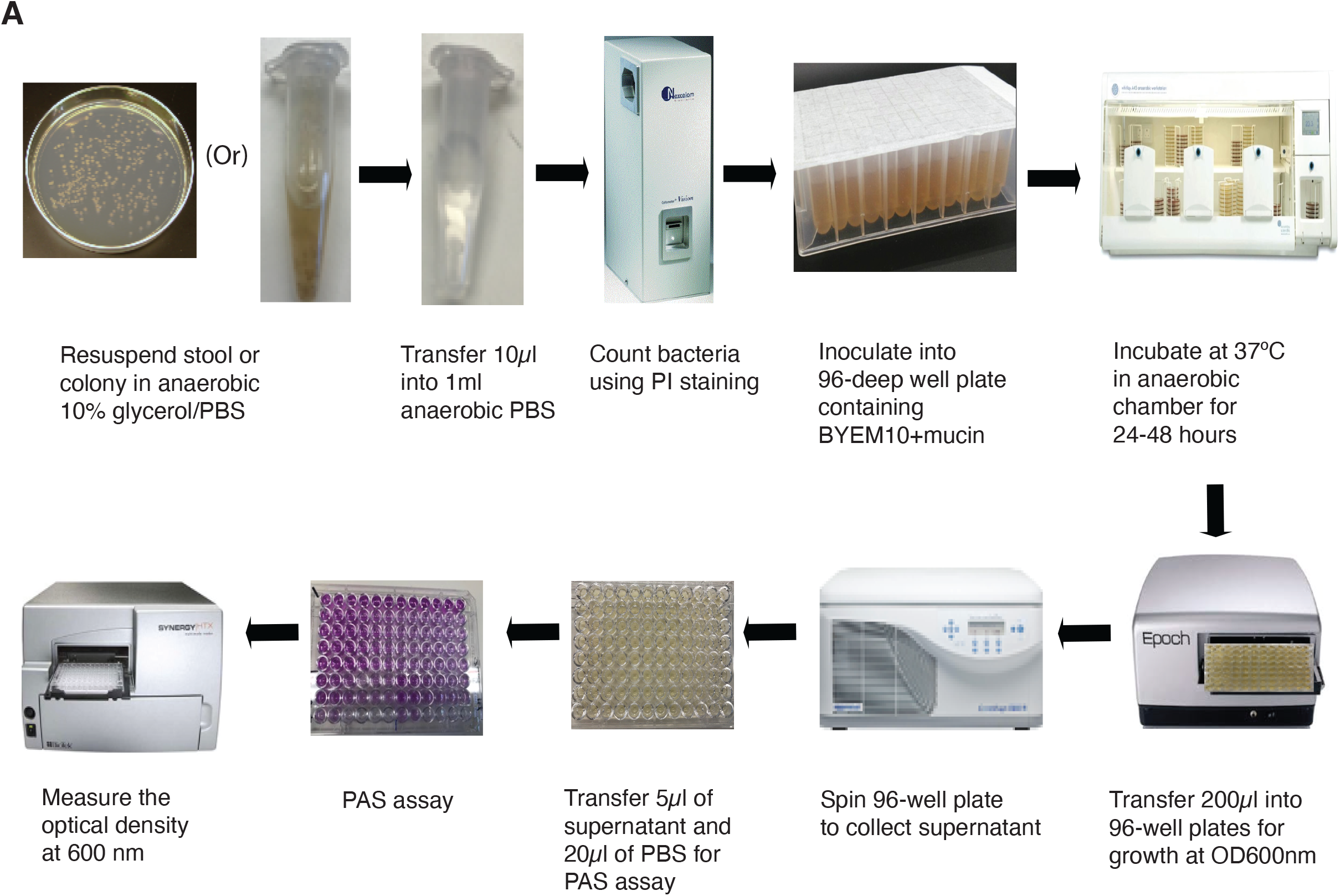
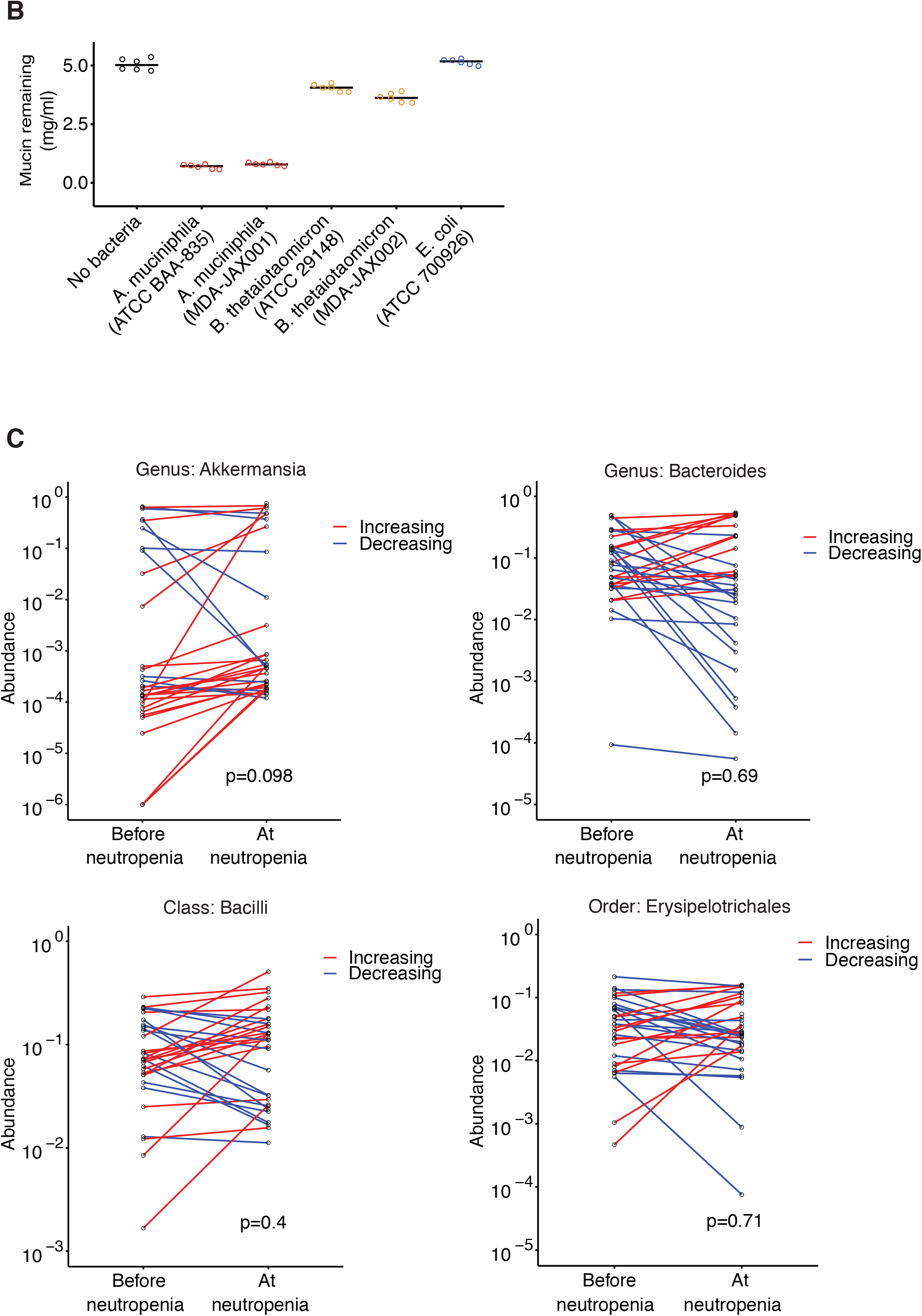
A) Workflow schematic of bacterial mucin glycan consumption assay. B) Results of mucin glycan quantification following 48-hour culture of indicated bacterial isolates. C) In the subset of patients who did not develop neutropenic fever, relative bacterial abundances of the indicated taxa in stool samples collected at onset of neutropenia were compared to results of a baseline stool sample collected earlier in the hospitalization, using the Wilcoxon signed-rank test.

**Figure S2.**
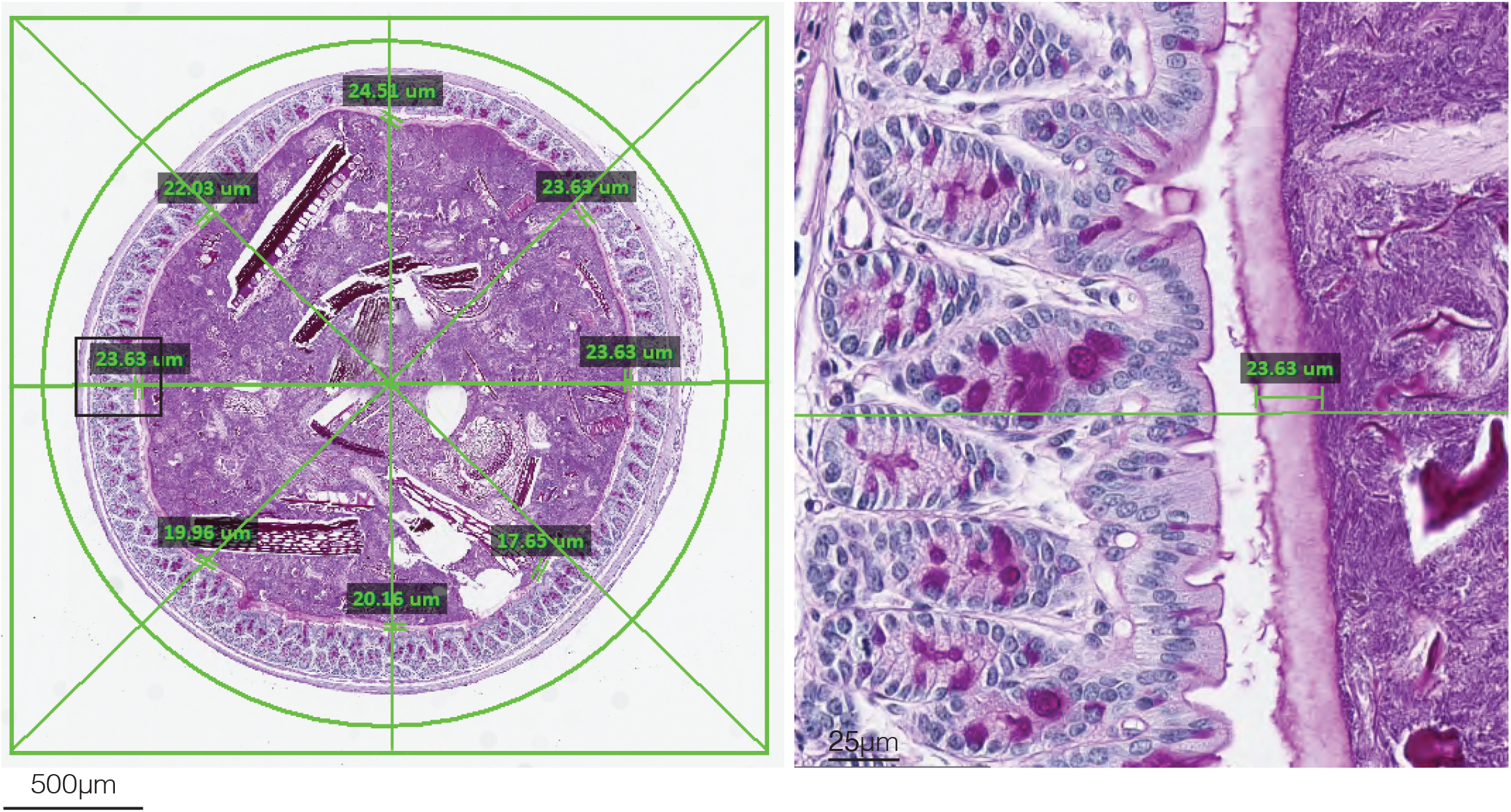
Schematic of histological quantification of the dense inner colonic mucus layer histologically, following PAS staining. Eight equally radially spaced sites are identified for mucus layer thickness quantification which are then averaged for each sample.

**Figure S3.**
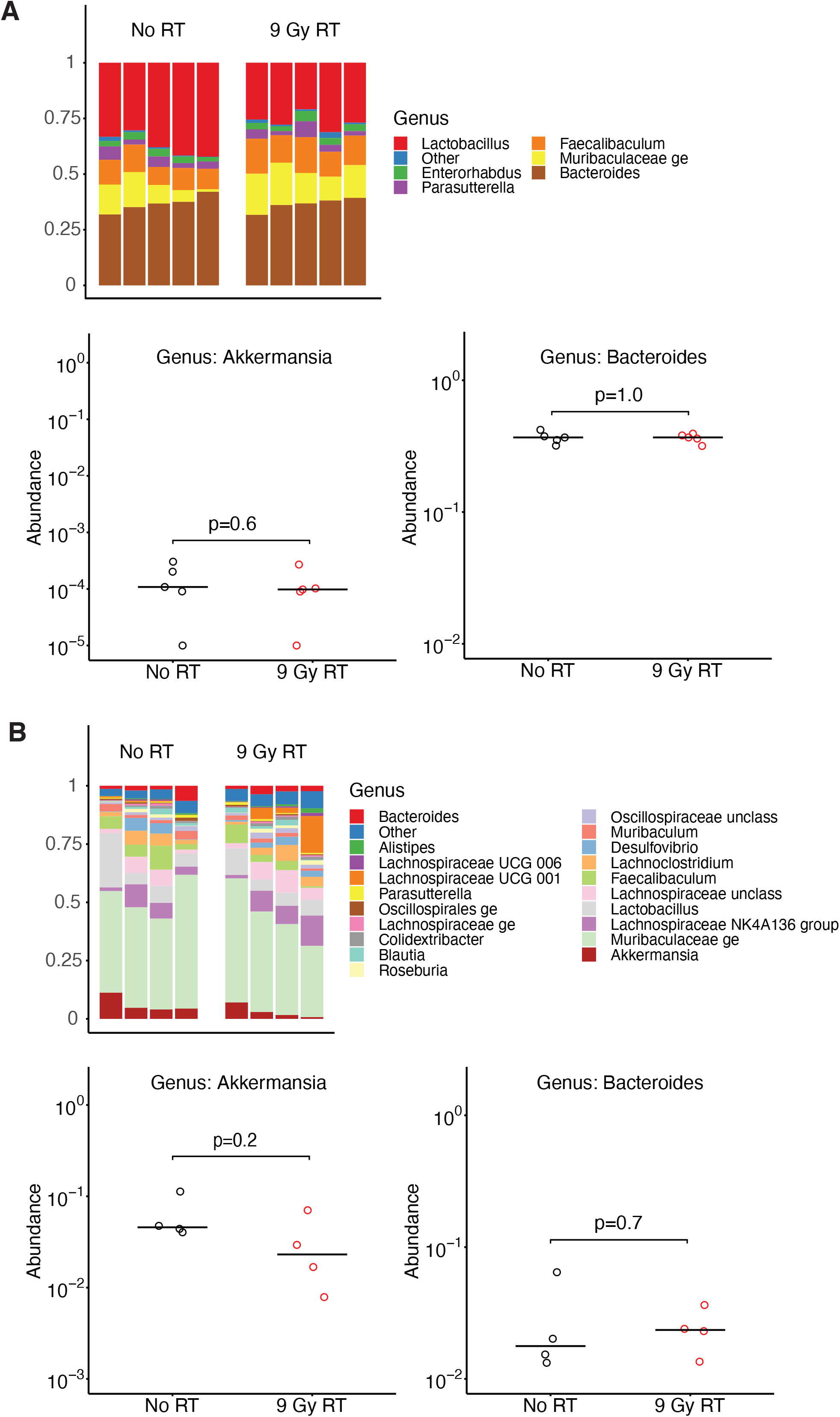
Radiation does not directly lead to a selective advantage for *Akkermansia* or *Bacteroides*. A) Fecal samples from normal mice were exposed to 9 Gy RT and then cultivated on Columbia blood agar plates in anaerobic conditions. Bacterial composition was determined by swabbing the plates and performing 16S rRNA gene sequencing. B) Fecal samples from normal mice were exposed to 9 Gy RT as in A), and then administered by gavage to mice following antibiotic decontamination with ampicillin, metronidazole and vancomycin in the drinking water. One week after fecal transplantation, stool pellets were collected and the bacterial composition was evaluated by 16S rRNA gene sequencing.

**Figure S4.**
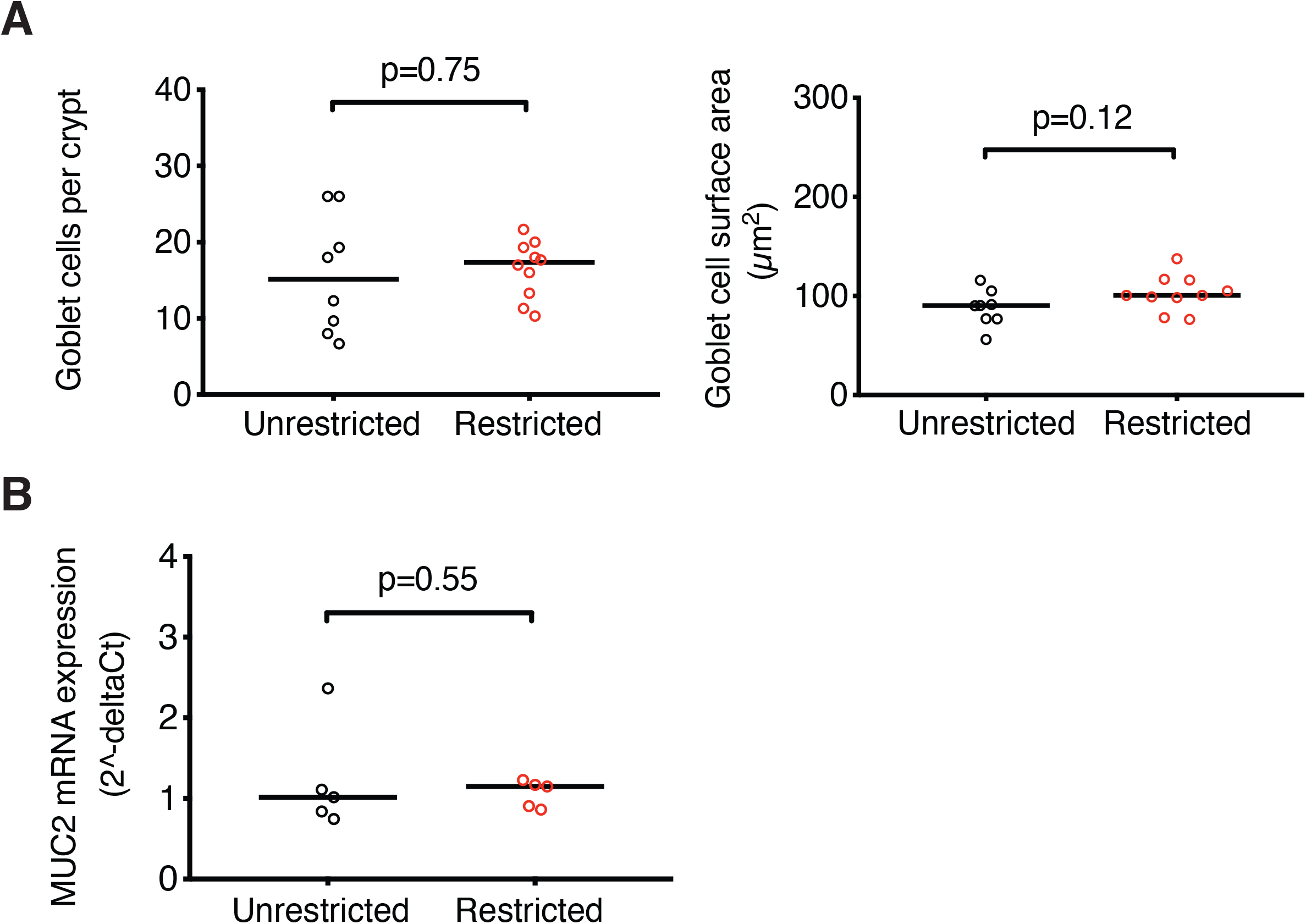
Dietary restriction has no clear impact on colonic mucus producing cells. A) Mice were subjected to dietary restriction for one week, and then colonic tissues were harvested and examined histologically. Goblet cell numbers were quantified, as well as goblet cell surface area. B) Gene expression of muc2 in colonic tissues was quantified in mice following one week of dietary restriction.

**Figure S5.**
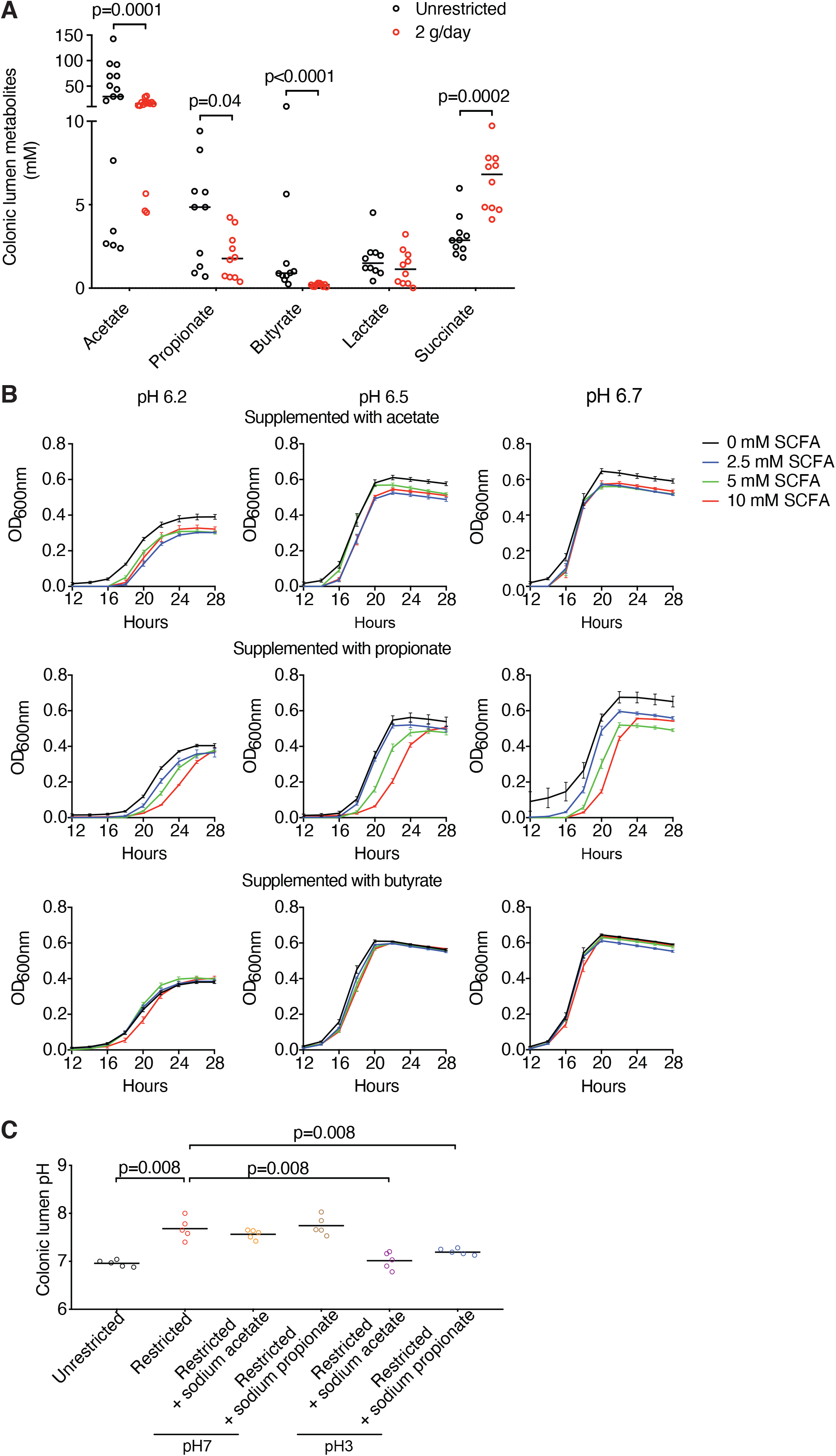
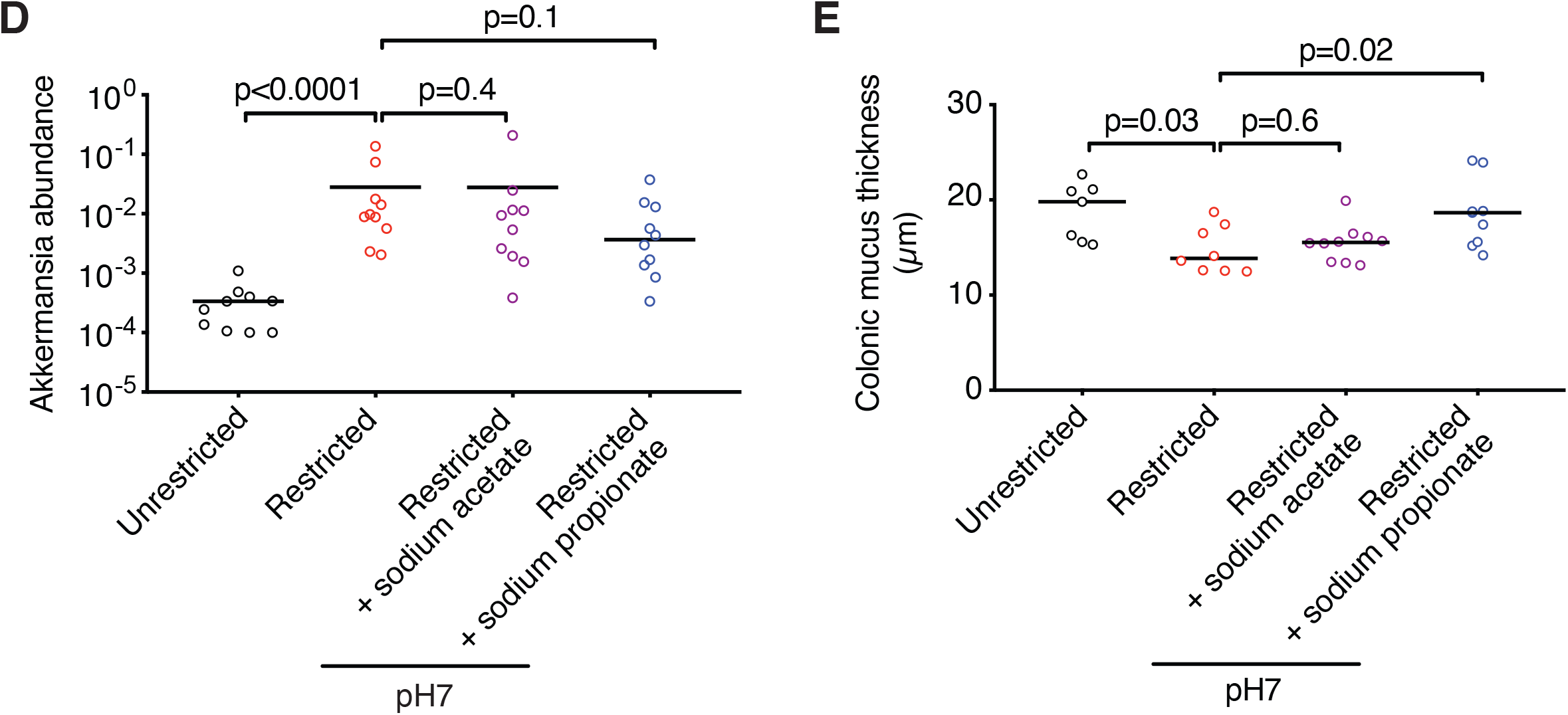
A) Raw values (without normalization) of metabolite quantification from sample depicted in Figure 4C. B) *A. muciniphila* growth in samples depicted in Figure 4E, quantified by optical density (OD) 600mm. C) Mice underwent dietary restriction and treatment with supplemental sodium acetate and sodium propionate adjusted to the indicated pH levels for one week, followed by quantification of the pH of colonic luminal contents. D) Mice were treated with dietary restriction and supplemental sodium acetate and sodium propionate adjusted to the indicated pH levels for one week, and fecal bacterial composition was evaluated by 16S rRNA gene sequencing; combined results of 2 experiments. E) Mice were treated as in D) and the colonic mucus thickness was quantified histologically; combined results of 2 experiments.

**Figure 6S.**
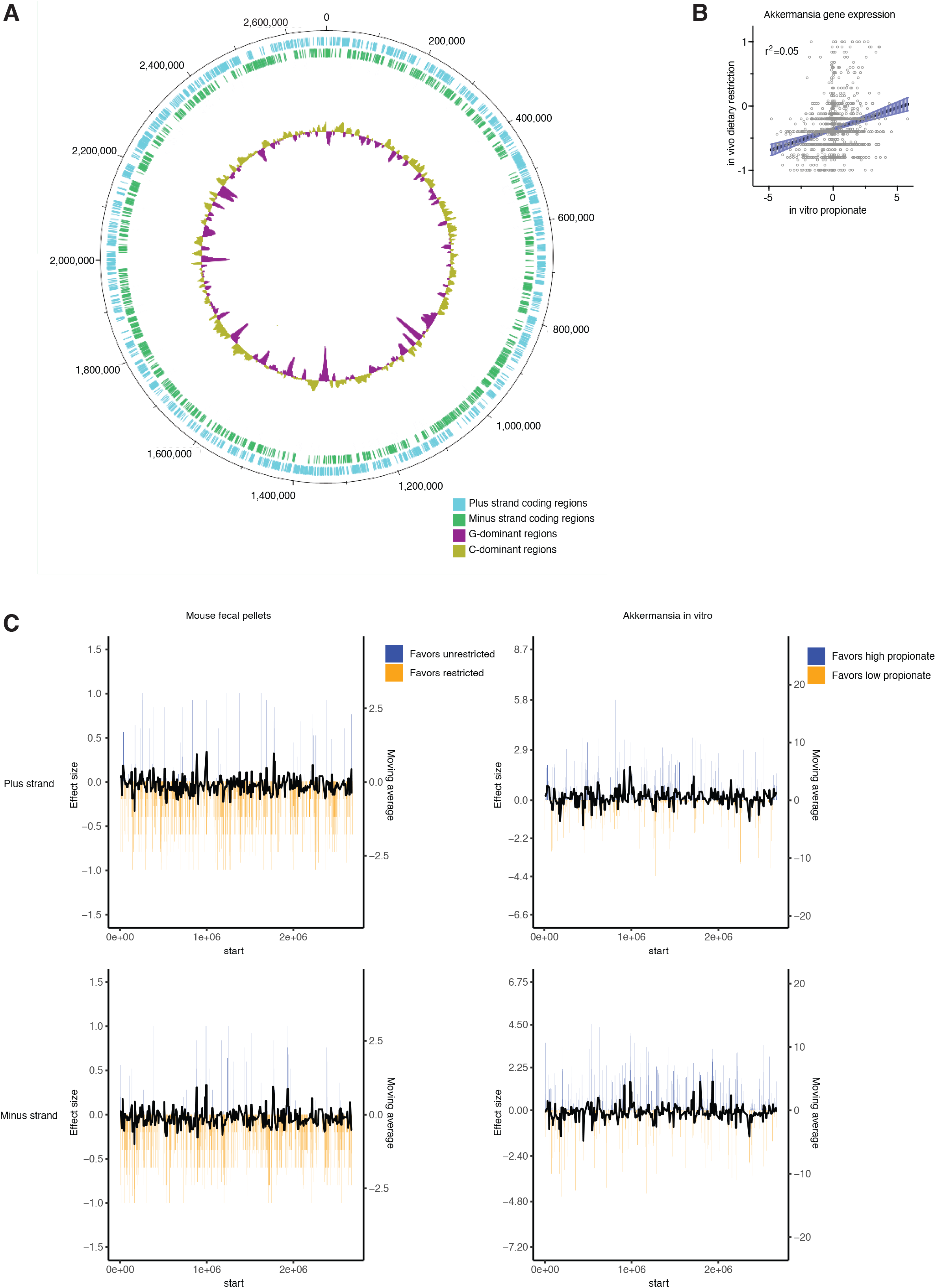
A) Circularized genome of *A. muciniphila* (MDA-JAX001). G- and C- dominant regions depict results of 10,000 bp moving averages. B) Changes in *A. muciniphila* gene expression were quantified in the settings of dietary restriction and in vitro exposure to propionate. Effect size statistics were quantified by the Mann-Whitney and Kruskal-Wallis methods, respectively, followed by using Spearman’s rank-order correlation test (p<0.0001). C) Changes in gene expression from a genomic perspective are depicted, along with 10,000 bp moving averages.

## Supplemental methods

### Human samples

Stool biospecimen collection from patients was approved by the University of Texas MD Anderson’s Institutional Review Board and signed informed consent was provided by all study participants. Samples were collected from patients undergoing stem cell transplantation and stored at 4°C for up to 48 hours before they were aliquoted for long-term storage at -80°C. Neutropenia was defined as a total white blood cell count less than 500 per microliter of blood. Neutropenic onset stool samples collected between day -2 to day +2 relative to the first day of neutropenia were eligible for inclusion in the study. Patients who received antibiotics other than standard bacterial prophylaxis with levofloxacin during the hospitalization prior to collection of neutropenic onset stool samples were excluded. Fever was defined as an oral temperature greater than 38.0°C within 4 days of neutropenic onset stool sample collection. For paired microbiome analyses, baseline stool samples were collected between 16 and 4 days (median 5.5 days) prior to neutropenic onset samples.

### Sequencing of 16S rRNA gene amplicons

Fecal samples were collected from patients and mice and weighed before DNA isolation. DNA was extracted from 20 mg to 200 mg of fecal material using the QIAamp Fast DNA Stool Mini Kit (Qiagen), following the manufacturer’s protocol, with additional heating and bead-beating steps. Methods for sequencing the V4 hypervariable region of the 16S rRNA gene were adapted from those developed for the Earth Microbiome Project (*1*). The quality and quantity of the barcoded amplicons were assessed on an Agilent 4200 TapeStation system (Agilent) and Qubit Fluorometer (Thermo Fisher Scientific), and libraries were prepared after pooling at equimolar ratios. The final libraries were sequenced with a 2×250 base pair paired-end protocol on the Illumina Miseq platform.

### Microbiome data analysis

Sequencing data from paired-end reads were de-multiplexed using QIIME (*2*). Merging of paired-end reads, dereplicating, and length filtering were performed using VSEARCH (*3*). Following de-noising and chimera calling using UNOISE3 (*4*), unique sequences were taxonomically classified with Mothur (*5*) using the Silva database (*6*) version 138. Weighted UniFrac distances (*7*) were determined using QIIME, visualized using principle coordinate analysis, and evaluated for statistical significance using Permutational Multivariate Analysis Of Variance (PERMANOVA) testing. For differential abundance analysis, abundances of sequences belonging to taxonomical groups at the genus, family, class and order levels were ranked in descending order for variance, maximum abundance, and median abundance, and the top 15 features for each criterion were included for analysis using the Mann-Whitney U test after logit transformation. P values were adjusted for multiple comparisons using the method of Benjamini and Hochberg. Paired samples were analyzed using the Wilcoxon signed rank test.

### Culturing of bacteria from stool samples

Stool samples from humans or mice were suspended in 1 ml of 10% anaerobic glycerol in a Whitley anaerobic chamber (10% H_2_, 5% CO_2_ and 85% N_2_) and quantified using a Nexcelom Cellometer cell counter with propidium iodide staining. Mucolytic bacteria including *Akkermansia muciniphila* (ATCC BAA-835), *Bacteroides thetaiotamicron* (ATCC 29148), *Akkermansia muciniphila* (novel murine isolate termed “MDA-JAX001”), *Bacteroides thetaiotamicron* (novel murine isolate termed “MDA-JAX002”) and a non-mucolytic *E. coli* (ATCC 700926) were used as controls. 1×10^6^ bacterial cells/ml was inoculated into 1.5 ml of BYEM10 broth (pH 7.2) with and without 5 mg/ml of porcine gastric mucin (PGM) (Sigma) and incubated up to 48 hours. Optical density (OD_600nm_) of the bacterial culture was measured with a BioTek EPOCH_2_ plate reader at 0, 24 and 48 hours of culture by transferring 200 µl of samples into flat bottom 96-well microtiter plates (Falcon). Culture supernatants were collected from each time point and stored at -30°C for quantification of post-culture levels of mucin glycans.

### Mucin consumption assay

Levels of mucin glycans in the culture supernatant were determined by a Periodic acid-Schiff assay as previously described (*8*) with minor modifications. Briefly, 5 µl of culture supernatant was transferred into a round bottom 96-well plate (Falcon) containing 20 µl of PBS. Serially diluted PGM standards were prepared, then 120 µl of freshly prepared 0.06% Periodic acid in 7% acetic acid was added, and incubated at 37°C for 90 minutes, followed by 100 µl of Schiff’s reagent (Sigma) and incubation at room temperature for 40 minutes. Absorbance was measured at 550 nm using a BioTek Synergy HTX plate reader.

### Mice

Studies in animal models conformed to the Guide for the Care and Use of Laboratory Animals Published by the US National Institutes of Health and was approved by the Institutional Animal Care and Use Committee. Six-to eight-week-old female C57BL/6 mice were obtained from the Jackson Laboratory.

### Total body radiotherapy

Mice were exposed to a single myeloablative dose of total body radiotherapy (9 Gy RT) using a Shepherd Mark I, Model 30, 137Cs irradiator.

### Histological methods

Colonic sections containing stool pellet were collected 1.5cm from the anus and immersed in fresh methanol-Carnoy’s fixative (methanol 60%, chloroform 30% and glacial acetic acid 10%) for 24 hours at room temperature as described previously (*9*). Preserved tissue was then incubated twice in dry methanol for 30 minutes each, followed by two incubations in absolute ethanol for 20 minutes each, two incubations in xylene for 25 minutes each, and 1 hour in paraffin at 60°C. Finally, the colon samples were embedded into paraffin blocks and 5 µm-thin sections were cut using the HistCore AUTOCUT microtome and placed on Superfrost Plus slides and dried overnight.

### Periodic acid-Schiff staining for mucus measurements

Colon sections were deparaffinized by incubating three times in xylene for 3 minutes each, two times for 2 minutes each in 100%, and 95% ethanol and washed with water three times for 1 minute each. PAS staining was performed using a Periodic acid-Schiff Staining Kit (Newcomer Supply). Briefly, tissue samples were incubated in 0.5% Periodic Acid for 10 minutes, washed 3 times in tap water for 5 minutes each and once in deionized water for 5 minutes. Samples were then incubated in Schiff’s Reagent for 20 minutes, washed with lukewarm tap water for 10 minutes, stained with hematoxylin for 5 minutes and washed in tap water for 2 minutes. Samples were quickly dipped in 1% acid alcohol, washed in tap water for 1 minutes and dipped 3-4 times in lithium carbonate. Finally, samples were rinsed three times in tap water for 5 minutes each followed by a 5-minute wash in deionized water and further dehydrated twice for 2 minutes each in 95% ethanol, three times for 2 minutes each in 100% ethanol and three times for 3 minutes each in xylene. Slides were mounted using Permount Mounting Media (SP15-100 Fisher Scientific) and sections were imaged using an Aperio AT2. Mucus thickness of the colonic sections were measured using eSlide Manager Version 12.4.3.5008. Eight measurements per image were taken and averaged over the entire usable colon surface. See Figure S2 for a representative image showing how measurement locations are divided and how the locations for quantifying the inner mucus layer are systematically identified.

### Melphalan therapy

Mice were administered a single subcutaneous injection of melphalan (20 mg/kg diluted in PBS). Control mice were injected with an equal volume of PBS.

### Irradiated murine fecal pellet culture

Mouse fecal pellets were suspended at 15 to 20 mg of stool per mL anaerobic PBS, strained through a 100 µm cell strainer, and exposed to 9 Gy or not. Fecal bacteria were plated on Columbia blood agar plates at a 10^−3^ dilution and grown in anaerobic conditions at 37°C. On day 7, bacterial colonies on each plate were collectively swabbed and evaluated by 16S sequencing.

### Fecal microbiome transplant

Mouse fecal pellets were prepared as described above, but instead of bacterial culture were introduced to mice by gavage (200 µL). Prior to fecal microbiome transplant, mice had been previously treated for 4 days with a decontaminating antibiotic cocktail (ampicillin: 1 mg/mL, metronidazole: 1 mg/mL, vancomycin: 1 mg/mL) administered in the drinking water, followed by a 2-day washout period. Stool samples were collected 7 days after fecal microbiome transplant for 16S sequencing.

### Metabolic monitoring

Tecniplast (Buguggiate, Italy) metabolic cages were used to determine food consumption in the radiation mouse model. The metabolic cages consist of a circular upper portion, which houses one mouse; a wire-grid floor; and a lower collection chamber with a specialized funnel that separates fecal pellets and urine. Pre-weighed food was introduced to each metabolic cage daily. Ground uneaten food was separated from the funnel containing stool and weighed. Daily food consumption was calculated by subtracting remaining chow in the hopper plus ground food from initial chow weight.

### Dietary restriction

Mice were subjected to an approximately 50% reduction of their normal oral intake for 7 days. Mice were provided daily with 2 grams of LabDiet PicoLab Rodent Diet 20 (5053), available in 1 and 5 gram tablets, per mouse on the floor of each cage with unlimited water.

### Narrow spectrum antibiotic administration

The dosage and concentrations of the antibiotics in the drinking water and gavage were based on published antibiotic doses and a projected daily consumption of 5 mL by an adult mouse. Antibiotics administered in the drinking water included azithromycin (0.35 mg/mL), vancomycin (0.625 mg/mL) and streptomycin (0.5 mg/mL) (*10, 11*). For the dietary restriction model, mice were pretreated with antibiotics in the drinking water beginning 5 days prior to the onset of dietary restriction and continued for 7 consecutive days thereafter. For the radiation mouse model, azithromycin (8.75 mg/mL in 200 µL) was administered by oral gavage once a day for 6 consecutive days after RT.

### Detection and quantification of goblet cells

Goblet cell surface areas and numbers were analyzed in periodic acid-Schiff stained distal colon sections described above. The number of goblet cells was counted in 3 crypts per mouse (average crypt length 150 µm) and in 6 to 10 mice per group. Goblet cell surface area, measured using ImageJ, was quantified in 20 cells per crypt, 3 crypts per mouse and 6 to 10 mice per group (a total of 360-600 Goblet cells per group).

### Quantitative PCR

Total RNA was extracted from homogenized tissue using the miRNeasy (Qiagen, Catalog: 217004). RNA was treated on column with DNAse to eliminate contaminating genomic DNA. The cDNA was synthesized using High Capacity cDNA Reverse Transcription Kit (Life tech, Catalog: 4368814). The mRNA levels of selected targets were quantified by qPCR using specific TaqMan probes (Muc2, Mm01276696_m1) and were normalized to GAPDH mRNA level (Gapdh, Mm99999915_g1).

### Bomb Calorimetry

Cecal luminal contents were dried for 12 hours using the Eppendorf Vacufuge, then weighed and heat of combustion was determined in a 6200 Isoperibol Calorimeter Equipped with a semi-micro oxygen combustion vessel. Benzoic acid was used as standard (Parr Instrument Company, Molline IL).

### pH measurement

Luminal stool isolated from the descending colon of mice was diluted with deionized water (5 µL water per mg stool). Samples were homogenized using a sterile wooden stick (Fisherbrand™ Plain-Tipped Applicators) and pH was measured at room temperature.

### Metabolite analysis

Metabolite standard compounds included the following (all compounds were from MilliporeSigma unless indicated otherwise): d_4_-acetic acid (Cambridge Isotope Laboratories); ^13^C_3_-lactic acid (Cambridge Isotope Laboratories), propionic acid, butyric acid, valeric acid, ketoleucine (4-methyl-2-oxovaleric acid), succinic acid, malic acid, glyceric acid, 2-hydroxyglutaric acid. Polar metabolites were extracted using water as described previously (*12*) with the following adjustments. Fecal samples were diluted 20-fold with deionized water based on sample weight and vortexed using a multi-tube vortexer (Fisher Scientific) for 20 minutes. Next, samples were centrifuged at 17,000 *g* for 10 min at 0°C, after which the supernatants were transferred to a 1.5 mL microfuge tube from which 50 µL was transferred to an LC-MS vial for Ion-Chromatography Mass-Spectrometry (IC-MS) analysis. The remaining supernatant was stored at -80°C. Samples were analyzed by IC-MS to broadly assess polar anionic metabolites as described previously (*13*) with the following modifications to ensure acquisition of low molecular weight short chain fatty acids (SCFAs). IC mobile phase A (MPA; weak) was water, and mobile phase B (MPB; strong) was 100 mM KOH. A Thermo Dionex ICS-5000+ IC system included an IonPac AS11-HC-4 µm (2 × 250 mm) column with column compartment kept at 30°C. The autosampler tray was chilled to 4°C. The mobile phase flow rate was 360 µL/minute, and the gradient elution program was: 0-10 minutes, 1% MPB; 10-24 minutes, 1-15% MPB; 24-40 minutes, 15-60% MPB; 40-44 minutes, 60-80% MPB; 44-48 minutes, 80-100% MPB; 48-58 minutes, 100% MPB; 58-58.1 minutes, 100-1% MPB; 58.1-60.5 minutes, 1% MPB. The total run time was 61 minutes. Data were acquired using a Thermo Orbitrap Fusion Tribrid Mass Spectrometer under ESI negative ionization mode at a resolution of 240,000 (FWHM at m/z 200) for MS1 acquisition (50–550 m/z). Sample injection volume was 10 µL. For absolute quantification, external standard curves were constructed using standards for acetic acid (d_4_-acetic acid), lactic acid (^13^C_3_-lactic acid), propionic acid, butyric acid, valeric acid, ketoleucine, succinic acid, malic acid, glyceric acid, and 2-hydroxyglutaric acid mixed at the following concentrations: 1, 2, 10, 50, 250, and 500 µM. All standards and metabolites were measured at an exact mass corresponding to proton loss, except for the standard for acetic acid. d_4_-Acetic acid rapidly exchanges a deuterium located at its carboxylic acid group with a hydrogen when dissolved in water, so acquisition m/z was set to proton loss for d_3_-acetic acid (m/z of 62.03252). Peak areas were integrated and exported to Excel using Thermo TraceFinder (version 5.0) software. Calibration curve construction and absolute quantification was performed in Excel (Microsoft).

### Effects of pH on *A. muciniphila*

The pH of the BYEM10 medium with 5mg/ml PGM was adjusted anaerobically. Each adjusted medium (1.5 ml) was dispensed into a 96-well deep-well plate and *A. muciniphila* (MDA-JAX001), or *E. coli* K-12 MG1655 was inoculated at 1×10^6^ cells/ml and incubated anaerobically up to 48 hours. Optical density (OD_600nm_) of the bacterial growth was measured at 48 hours and culture supernatants were collected and stored at -30°C for post-culture quantification of mucin glycans (described above).

### Effects of short chain fatty acids on *A. muciniphila*

To evaluate the effects of SCFAs on mucolytic activity, 1×10^4^ *A. muciniphila* (MDA-JAX001) was inoculated into 1.5 ml of BYEM10 (pH6.5) with porcine gastric mucin together with varying concentrations of sodium propionate (2.5 mM, 5 mM, 10 mM and 20 mM), sodium butyrate (2.5 mM, 5 mM, 10 mM), or sodium acetate (2.5 mM, 5 mM, 10 mM, 20 mM) (*14*). ***Growth kinetics***. 250 µl of inoculated medium was transferred into 96-well cell culture plate where OD_600nm_ was read automatically every 2 hours for 48 hours. ***Mucus glycan consumption***. After 24h, 200 µl of culture suspension was centrifuged at 4000g at 4°C for 15 minutes. Supernatant was collected for post-culture quantification of mucin glycans.

### Short-chain fatty acid (SCFA) administration

Sodium propionate (Sigma, 150 mM) and sodium acetate (Sigma, 150 mM) were administered as reported previously (*14*). For the dietary restriction model, SCFA were introduced in the drinking water upon the onset of calorie restriction and continued for 7 consecutive days thereafter. For the radiation mouse model, sodium propionate was introduced in the drinking water beginning 5 days prior to RT to avoid further impacting on water consumption which is compromised in mice following RT. In addition to drinking water supplementation, sodium propionate was administered by oral gavage (200 µl, 300 mM) twice a day after RT.

### Murine temperature monitoring

Ocular temperatures were measured using an infrared FLIR E60 camera (FLIR, UK) as previously described (*15*). Briefly, a 20mm lens was attached to the front of the camera using a 3D printed lens holder without any modifications to the camera for close-up imaging. Ocular temperatures were acquired with a 56 mm focal distance perpendicular to the eye being assessed. Data were analyzed by FLIR Tools+ software.

### Murine colonic tissue cytokine analysis

250 mg of colonic tissue per mL of extraction buffer (Thermofisher Scientific) was homogenized and centrifuged for 5 minutes at 10000g. Supernatant was used from each sample to determine cytokine levels. The ProcartaPlex Multiplex immunoassay (Mouse monitoring 48-plex Ref. EPX480-20834-901 and Mouse custom ProcartaPlex 15-plex) was conducted per the manufacturer’s instructions (Affymetrix). Results were acquired with a MagPix instrument and analyzed with xPONENT software (Luminex Corporation).

### Whole genome sequencing of *Akkermansia muciniphila* (MDA-JAX001)

*A. muciniphila* (MDA-JAX001) genomic DNA was isolated and purified using Qiagen Genomic-tip 20/G, according to the manufacturer’s instructions. For long-read Nanopore sequencing, 400 ng of genomic DNA was used for library preparation using the Rapid Sequencing Kit (SQK-RAD004, Oxford Nanopore Technologies). The library was loaded into a FLO-MIN106 flow-cell for a 12h sequencing run on a MinION sequencer platform (Oxford Nanopore Technologies, Oxford, UK). Data acquisition and real-time base calling were carried out by the MinKNOW software. The fastq files were generated from the base called sequencing fast5 reads. For short-read illumina sequencing, libraries were constructed using Nextera DNA Flex Library Prep Kit (Illumina), following manufacturer’s protocol. The final libraries were loaded into the NovaSeq 6000 platform (Illumina) and sequenced 2×150 bp paired-end read, resulting in ∼5 Gb per sample. To assemble the complete genome information of *A. muciniphila* (MDA-JAX001), we used the SPAdes software (ver 3.13.1.) with the long and short reads combined (*16*). After obtaining a de-Bruijn graph, we confirmed a circular structure in the longest scaffold by DNA plotter software (ver 4.44.1) within the first and last 460 bps (*17*). The genome was annotated using Prokka to identify putative coding sequences (*18*) followed by alignment to the CAZy database (*19*) available in the dbCAN2 server (*20*), and to the NCBI RefSeq protein database maintained by the National Center for Biotechnology Information, using DIAMOND (*21*).

### Bacterial RNA sequencing

#### In vitro A. muciniphila

*A. muciniphila* (MDA-JAX001) was grown in BYEM10 (pH 6.5) with porcine gastric mucin (5mg/ml) broth and varying concentrations of sodium propionate. When the samples without supplemental propionate reached an OD600nm of ∼0.3, corresponding to the exponential growth phase, all samples were harvested for total RNA isolation using an RNeasy Mini Kit (Qiagen, Valencia, CA). RNA was treated on column with DNAse to eliminate contaminating genomic DNA. RNA quality and quantity was determined using the TapeStation 4200 (Agilent Technologies, Santa Clara, CA).

#### RNA isolation from mouse fecal pellets

Approximately 30 mg of stool was freshly collected in 700 µL of ice cold Qiazol containing 200 µL of 0.1 mm diameter Zirconia Silica beads (BioSpec Cat. No. 11079101z). Samples were bead beaten twice for 2 min with a 30 second recovery in between. Samples were then centrifuged at 12,000 g for 1 min and supernatant was collected for RNA isolation using the miRNeasy mini kit (Qiagen 217004). RNA was treated on column with DNAse to eliminate contaminating genomic DNA. RNA quantity and quality was determined using the TapeStation 4200.

#### RNA sequencing and analysis

250 ng of total RNA from mouse stool or bacterial cultures was used to construct libraries using the Nugen Ovation Complete Prokaryotic RNA-Seq Systems (NuGen), following manufacturer’s protocol. The cDNA libraries were sequenced on the Illumina MiSeq system for 1×300 bp single-read run, resulting in ∼1 million reads per sample (Illumina, San Diego, CA, USA).

Sequence data were split using QIIME (version 1.9.1 and their qualities were checked by VSEARCH (version 2.14.1) (*2, 3*). Data were filtered and truncated on their quality by VSEARCH default settings. Adapter sequences were removed using QIIME. The total reads per mouse stool sample were 1969892 ± 562811 (mean ± standard deviation) and the total reads from bacterial cultures were 310963 ± 67017 (mean ± standard deviation).

Sequences of ribosomal RNA were removed using BWA (ver 0.7.17) and the prokaryotic ribosomal RNA sequences in the NCBI RefSeq prokaryotic genome database (*22, 23*). *A. muciniphila* MDA-JAX001 sequences were then identified using DIAMOND to map to the results of our whole genome sequencing described above as a reference. Associations between gene expression and varying propionate concentrations in vitro were quantified using the Kruskal-Wallis rank sum test after logit transformation, while associations with dietary restriction were quantified using the Mann-Whitney U test. Given the exploratory nature of these metatranscriptomic analyses, p values have not been corrected for multiple comparisons.

## References

1. N. M. Kuderer, D. C. Dale, J. Crawford, L. E. Cosler, G. H. Lyman, Mortality, morbidity, and cost associated with febrile neutropenia in adult cancer patients. Cancer 106, 2258–2266 (2006).

2. E. Tai, G. P. Guy, A. Dunbar, L. C. Richardson, Cost of Cancer-Related Neutropenia or Fever Hospitalizations, United States, 2012. J Oncol Pract 13, e552–e561 (2017).

3. R. A. Taplitz et al., Antimicrobial Prophylaxis for Adult Patients With Cancer-Related Immunosuppression: ASCO and IDSA Clinical Practice Guideline Update. J Clin Oncol 36, 3043–3054 (2018).

4. G. P. Bodey, The changing face of febrile neutropenia-from monotherapy to moulds to mucositis. Fever and neutropenia: the early years. J Antimicrob Chemother 63 Suppl 1, i3–13 (2009).

5. A. G. Freifeld et al., Clinical practice guideline for the use of antimicrobial agents in neutropenic patients with cancer: 2010 update by the infectious diseases society of america. Clin Infect Dis 52, e56–93 (2011).

6. F. B. Tamburini et al., Precision identification of diverse bloodstream pathogens in the gut microbiome. Nat Med 24, 1809–1814 (2018).

7. Y. Taur et al., Intestinal Domination and the Risk of Bacteremia in Patients Undergoing Allogeneic Hematopoietic Stem Cell Transplantation. Clin Infect Dis, (2012).

8. J. F. Sicard, G. Le Bihan, P. Vogeleer, M. Jacques, J. Harel, Interactions of Intestinal Bacteria with Components of the Intestinal Mucus. Front Cell Infect Microbiol 7, 387 (2017).

9. M. Kilcoyne, J. Q. Gerlach, M. P. Farrell, V. P. Bhavanandan, L. Joshi, Periodic acid-Schiff’s reagent assay for carbohydrates in a microtiter plate format. Anal Biochem 416, 18–26 (2011).

10. I. Lagkouvardos et al., Sequence and cultivation study of Muribaculaceae reveals novel species, host preference, and functional potential of this yet undescribed family. Microbiome 7, 28 (2019).

11. B. Gyurkocza, B. M. Sandmaier, Conditioning regimens for hematopoietic cell transplantation: one size does not fit all. Blood 124, 344–353 (2014).

12. E. Ansaldo et al., Akkermansia muciniphila induces intestinal adaptive immune responses during homeostasis. Science 364, 1179–1184 (2019).

13. J. M. Macy, L. G. Ljungdahl, G. Gottschalk, Pathway of succinate and propionate formation in Bacteroides fragilis. J Bacteriol 134, 84–91 (1978).

14. Q. P. Liu et al., Bacterial glycosidases for the production of universal red blood cells. Nat Biotechnol 25, 454–464 (2007).

15. E. H. Crost et al., The mucin-degradation strategy of Ruminococcus gnavus: The importance of intramolecular trans-sialidases. Gut Microbes 7, 302–312 (2016).

16. B. Vogel et al., Touch-free measurement of body temperature using close-up thermography of the ocular surface. MethodsX 3, 407–416 (2016).

17. A. A. Romanovsky et al., Fever and hypothermia in systemic inflammation: recent discoveries and revisions. Front Biosci 10, 2193–2216 (2005).

18. W. Tao, D. J. Deyo, D. L. Traber, W. E. Johnston, E. R. Sherwood, Hemodynamic and cardiac contractile function during sepsis caused by cecal ligation and puncture in mice. Shock 21, 31–37 (2004).

19. C. W. Png et al., Mucolytic bacteria with increased prevalence in IBD mucosa augment in vitro utilization of mucin by other bacteria. The American journal of gastroenterology 105, 2420–2428 (2010).

20. J. Sun et al., Therapeutic Potential to Modify the Mucus Barrier in Inflammatory Bowel Disease. Nutrients 8, (2016).

21. Y. Shono et al., Increased GVHD-related mortality with broad-spectrum antibiotic use after allogeneic hematopoietic stem cell transplantation in human patients and mice. Science translational medicine 8, 339ra371 (2016).

22. C. M. Dejea et al., Patients with familial adenomatous polyposis harbor colonic biofilms containing tumorigenic bacteria. Science 359, 592–597 (2018).

23. M. Derrien, E. E. Vaughan, C. M. Plugge, W. M. de Vos, Akkermansia muciniphila gen. nov., sp. nov., a human intestinal mucin-degrading bacterium. Int J Syst Evol Microbiol 54, 1469–1476 (2004).

24. S. S. Seregin et al., NLRP6 Protects Il10-/-Mice from Colitis by Limiting Colonization of Akkermansia muciniphila. Cell Rep 19, 733–745 (2017).

25. S. M. Bloom et al., Commensal Bacteroides species induce colitis in host-genotype-specific fashion in a mouse model of inflammatory bowel disease. Cell Host Microbe 9, 390–403 (2011).

26. N. A. Pudlo et al., Symbiotic Human Gut Bacteria with Variable Metabolic Priorities for Host Mucosal Glycans. mBio 6, e01282–01215 (2015).

27. A. Bell et al., Elucidation of a sialic acid metabolism pathway in mucus-foraging Ruminococcus gnavus unravels mechanisms of bacterial adaptation to the gut. Nat Microbiol 4, 2393–2404 (2019).

28. A. B. Hall et al., A novel Ruminococcus gnavus clade enriched in inflammatory bowel disease patients. Genome Med 9, 103 (2017).

29. C. Depommier et al., Supplementation with Akkermansia muciniphila in overweight and obese human volunteers: a proof-of-concept exploratory study. Nat Med 25, 1096–1103 (2019).

30. B. Routy et al., Gut microbiome influences efficacy of PD-1-based immunotherapy against epithelial tumors. Science 359, 91–97 (2018).

31. C. Ozkul, M. Yalinay, T. Karakan, Islamic fasting leads to an increased abundance of Akkermansia muciniphila and Bacteroides fragilis group: A preliminary study on intermittent fasting. Turk J Gastroenterol 30, 1030–1035 (2019).

32. A. Basolo et al., Effects of underfeeding and oral vancomycin on gut microbiome and nutrient absorption in humans. Nat Med 26, 589–598 (2020).

33. L. Li et al., The effects of daily fasting hours on shaping gut microbiota in mice. BMC Microbiol 20, 65 (2020).

34. E. A. Miyasaka et al., Total parenteral nutrition-associated lamina propria inflammation in mice is mediated by a MyD88-dependent mechanism. J Immunol 190, 6607–6615 (2013).

35. M. S. Desai et al., A Dietary Fiber-Deprived Gut Microbiota Degrades the Colonic Mucus Barrier and Enhances Pathogen Susceptibility. Cell 167, 1339–1353 e1321 (2016).

36. S. Jeong, H. Y. Kim, A. R. Kim, C. H. Yun, S. H. Han, Propionate Ameliorates Staphylococcus aureus Skin Infection by Attenuating Bacterial Growth. Front Microbiol 10, 1363 (2019).

37. L. W. Chia et al., Deciphering the trophic interaction between Akkermansia muciniphila and the butyrogenic gut commensal Anaerostipes caccae using a metatranscriptomic approach. Antonie Van Leeuwenhoek 111, 859–873 (2018).

38. M. T. Sorbara et al., Inhibiting antibiotic-resistant Enterobacteriaceae by microbiota-mediated intracellular acidification. J Exp Med 216, 84–98 (2019).

## References

1. L. R. Thompson et al., A communal catalogue reveals Earth’s multiscale microbial diversity. Nature 551, 457–463 (2017).

2. J. G. Caporaso et al., QIIME allows analysis of high-throughput community sequencing data. Nat Methods 7, 335–336 (2010).

3. T. Rognes, T. Flouri, B. Nichols, C. Quince, F. Mahe, VSEARCH: a versatile open source tool for metagenomics. PeerJ 4, e2584 (2016).

4. R. C. Edgar, UNOISE2: improved error-correction for Illumina 16S and ITS amplicon sequencing. bioRxiv, 081257 (2016).

5. P. D. Schloss et al., Introducing mothur: open-source, platform-independent, community-supported software for describing and comparing microbial communities. Appl Environ Microbiol 75, 7537–7541 (2009).

6. C. Quast et al., The SILVA ribosomal RNA gene database project: improved data processing and web-based tools. Nucleic Acids Res 41, D590–596 (2013).

7. C. Lozupone, M. E. Lladser, D. Knights, J. Stombaugh, R. Knight, UniFrac: an effective distance metric for microbial community comparison. ISME J 5, 169–172 (2011).

8. M. Kilcoyne, J. Q. Gerlach, M. P. Farrell, V. P. Bhavanandan, L. Joshi, Periodic acid-Schiff’s reagent assay for carbohydrates in a microtiter plate format. Anal Biochem 416, 18–26 (2011).

9. M. E. Johansson, G. C. Hansson, Preservation of mucus in histological sections, immunostaining of mucins in fixed tissue, and localization of bacteria with FISH. Methods Mol Biol 842, 229–235 (2012).

10. R. Li et al., Effects of oral florfenicol and azithromycin on gut microbiota and adipogenesis in mice. PLoS One 12, e0181690 (2017).

11. A. M. Schubert, H. Sinani, P. D. Schloss, Antibiotic-Induced Alterations of the Murine Gut Microbiota and Subsequent Effects on Colonization Resistance against Clostridium difficile. mBio 6, e00974 (2015).

12. S. Moosmang et al., Metabolomic analysis-Addressing NMR and LC-MS related problems in human feces sample preparation. Clin Chim Acta 489, 169–176 (2019).

13. Y. Sun et al., Functional Genomics Reveals Synthetic Lethality between Phosphogluconate Dehydrogenase and Oxidative Phosphorylation. Cell Rep 26, 469–482 e465 (2019).

14. P. M. Smith et al., The microbial metabolites, short-chain fatty acids, regulate colonic Treg cell homeostasis. Science 341, 569–573 (2013).

15. B. Vogel et al., Touch-free measurement of body temperature using close-up thermography of the ocular surface. MethodsX 3, 407–416 (2016).

16. A. Bankevich et al., SPAdes: a new genome assembly algorithm and its applications to single-cell sequencing. J Comput Biol 19, 455–477 (2012).

17. E. L. Sonnhammer, R. Durbin, A dot-matrix program with dynamic threshold control suited for genomic DNA and protein sequence analysis. Gene 167, GC1–10 (1995).

18. T. Seemann, Prokka: rapid prokaryotic genome annotation. Bioinformatics 30, 2068–2069 (2014).

19. V. Lombard, H. Golaconda Ramulu, E. Drula, P. M. Coutinho, B. Henrissat, The carbohydrate-active enzymes database (CAZy) in 2013. Nucleic Acids Res 42, D490–495 (2014).

20. H. Zhang et al., dbCAN2: a meta server for automated carbohydrate-active enzyme annotation. Nucleic Acids Res 46, W95–W101 (2018).

21. B. Buchfink, C. Xie, D. H. Huson, Fast and sensitive protein alignment using DIAMOND. Nat Methods 12, 59–60 (2015).

22. H. Li, R. Durbin, Fast and accurate short read alignment with Burrows-Wheeler transform. Bioinformatics 25, 1754–1760 (2009).

23. T. Tatusova et al., NCBI prokaryotic genome annotation pipeline. Nucleic Acids Res 44, 6614–6624 (2016).

